# Challenges and pitfalls of inferring microbial growth rates from lab cultures

**DOI:** 10.1101/2022.06.24.497412

**Authors:** Ana-Hermina Ghenu, Loïc Marrec, Claudia Bank

**Affiliations:** Instituto Gulbenkian de Ciência, Rua da Quinta Grande 6, Oeiras, 2780-156, Portugal; Institut für Ökologie und Evolution, Universität Bern, Baltzerstrasse 6, CH-3012 Bern, Switzerland; Swiss Institute of Bioinformatics, 1015 Lausanne, Switzerland

## Abstract

After more than 100 years of generating monoculture batch culture growth curves, microbial ecologists and evolutionary biologists still lack a reference method for inferring growth rates. Our work highlights the challenges of estimating the growth rate from growth curve data and shows that inaccurate estimates of growth rates significantly impact the estimated relative fitness, a principal quantity in evolution and ecology. First, we conducted a literature review and found which different types of methods are currently used to estimate growth rates. These methods differ in the meaning of the estimated growth rate parameter. Kinetic models estimate the intrinsic growth rate *µ* whereas statistical methods – both model-based and model-free – estimate the maximum *per capita* growth rate *µ*_max_. Using math and simulations, we show the conditions in which *µ*_max_ is not a good estimator of *µ*. Then, we demonstrate that inaccurate absolute estimates of *µ* is not overcome by calculating relative values. Importantly, we find that poor approximations for *µ* sometimes lead to wrongly classifying a beneficial mutant as deleterious. Finally, we re-analyzed four published data-sets using most of the methods found by our literature review. We detected no single best-fitting model across all experiments within a data-set and found that the Gompertz models, which were among the most commonly used, were often among the worst fitting. Our study provides suggestions for how experimenters can improve their growth rate and associated relative fitness estimates and highlights a neglected but fundamental problem for nearly everyone who studies microbial populations in the lab.

## 1 Introduction

Measuring batch culture growth curves is a common method used by nearly all who work with single-celled organisms in the laboratory. Growth curves allow experimenters to readily measure population phenotypes, like the dynamics and efficiency of growth in particular environments, for microscopic organisms whose individual cell phenotypes are often laborious or expensive to quantify. The growth rate is an especially important trait for evolutionary microbiologists and microbial ecologists. The growth rate is important because it is related to fitness in population biology, it is used to estimate the number of generations a microbial culture has been growing for (e.g., Wein and Dagan 2019), it is more responsive to selection that other traits in microbial evolution experiments (Wahl and Zhu, 2015), and it is central in describing competition for limited resources (Miller et al., 2005; Bernhardt et al., 2020). Overall, growth curves are commonly used because they are easy to obtain, have been used for a long time, and usually give consistent results within an experiment. The importance of growth curves is only increasing in the age of high-throughput experimental screens of microbial populations, from which conclusions are drawn about responses to ecological challenges, to antimicrobial drugs, and about optimal strains for agricultural purposes. Nevertheless, despite the popularity of gathering growth curve data and the proliferation of methods for extracting growth parameters from said data, it is not clear what is the best method for estimating values of interest for these data.

The idea behind the batch culture growth curve is simple: inoculate a sterile culture medium with a small number of individuals *N*_0_ and track the increase in population size over time using any available method to estimate population size (e.g., colony forming units, optical density, microscopy cells counts, flow cytometry). An idealized growth curve is shown in Figure 1, in which each panel shows the same simulated data but with a different y-axis. When an experimenter assesses a growth curve, what they may first observe is perhaps a “lag phase” with little growth. Then, they will *always* observe a phase of rapid growth, alternately called the “log phase” or “linear phase” by different researchers, in which the growth is linear when shown with the y-axis on a log-scale (Figure 1B). Figure 1C shows the first derivative of Figure 1B: in other words, the instantaneous *per capita* growth rate. The fastest growth rate is usually reached during the “log/linear phase” (Figure 1C; Monod 1949). Then, this phase may be followed by more decelerations and subsequent accelerations under diauxic/diphasic growth conditions (not shown or further discussed herein). Eventually the population growth slows down to halt at the “stationary phase”, reaching the final carrying capacity (denoted *K* in linear-scale, Figure 1A, or *A* in log-scale, Figure 1B) when all the usable resources are depleted from the batch culture.

**Figure 1:**
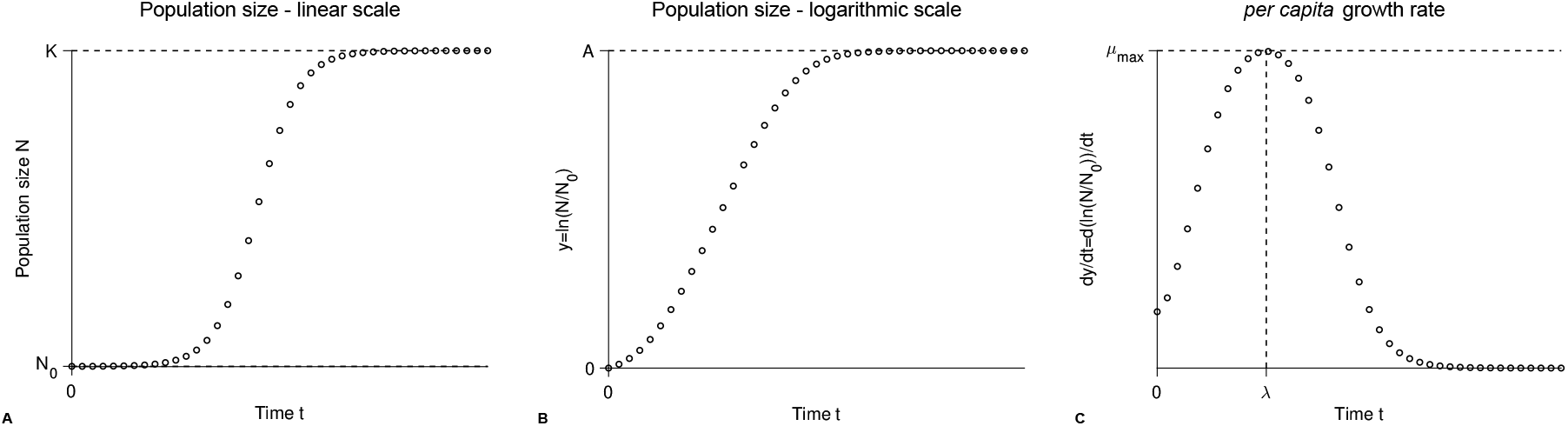
A schematic illustration of the same simulated batch culture population growth curve, plotted in three different ways.: **A)** Population size *N* versus time *t*. The quantity *K* corresponds to the carrying capacity and *N*_0_ to the initial population size. The initial population density or fraction is given by *N*_0_*/K*. **B)** Logarithm of the population size divided by the initial population size *y* = ln(*N/N*_0_) versus time *t*. Note that in other publications the quantity ln(*N/N*_0_) is sometimes denoted as *y*. The quantity *A* = ln(*K/N*_0_) is the logarithm of the fold increase over the initial population size at carrying capacity. **C)** First derivative of the logarithm of the population size divided by the initial population size d*y/*d*t* = d ln(*N/N*_0_)*/*d*t* versus time *t*. The function d*y/*d*t* may be interpreted as the *per capita* growth rate. The quantity *µ*_max_ is the *per capita* maximum growth rate and *λ* the lag phase duration.

Growth curves are often used to estimate the *per capita* growth rate and fitness. The *per capita* growth rate is important in population biology because it is used to calculate the growth of a mutant strain as compared to a wild-type strain: this is the relative growth rate, or the relative fitness (*w*). The relative fitness is particularly important in evolutionary biology since it classifies a mutant as deleterious (*w <* 1), neutral (*w* = 0) or beneficial (*w >* 1) with respect to natural selection. Indeed, when the relative fitness is greater than 1, the mutant reproduces faster than the wild-type, and conversely when *w* is less than 1. Although there is still discussion in the field about whether the relative growth rates measured from monoculture growth curves are predictive of competitive fitness (Concepción-Acevedo et al., 2015; Ram et al., 2019), many biologists use the growth rate as a measure of fitness (e.g., Knopp and Andersson 2018).

Microbial batch culture protocols have been used for over 100 years in microbiology (e.g., Slator 1916) and population ecology (e.g., Carlson 1913), and remain a mainstay of experimental evolution and ecology. During this time, many experimental protocols (Delaney et al., 2013; Hall et al., 2014; Stevenson et al., 2016; Kurokawa and Ying, 2017) and estimation methods (Zwietering et al., 1990; Baranyi and Roberts, 1994; Jung et al., 2015) have been developed for this type of data. Since the early 1990s, automated plate readers that incubate and periodically scan the opacity of the cultures growing in the microwells have simplified the process of gathering data for hundreds of bacterial populations simultaneously growing in (relatively) homogeneous batch culture environments. Sources of inconsistency, like the batch effect (Blomberg, 2011), can be mitigated, for example by growing all cultures of interest on the same day(s), in order to arrive at consistent data. Nevertheless, despite the long tradition and good recommendations for setting up experiments, programs and papers detailing methods for estimating growth rates (and other growth parameters) from this data continue to be published and highly sought after. Many of these estimation methods are implementations of classical models (e.g. Sprouffske and Wagner 2016; Petzoldt 2020). This shows that also after 100+ years of generating growth curve data, microbial ecologists and evolutionary biologists are still struggling to find the best way of estimating the growth rate from their data.

The main goal of our paper is to demonstrate that there are significant limitations to existing methods for using batch culture growth curve data to estimate the intrinsic growth rate *µ*, which is the fastest *per capita* number of divisions per time unit possible when the cell’s resources are limitless or otherwise optimal. These limitations impact the calculation of quantities of interest such as the selection coefficient and the relative fitness. We first take stock of how the community currently analyzes growth curve data by semi-quantitatively reviewing the literature to survey which methods are used in evolution and ecology. After explaining different approaches for modelling growth curves, we then use math and simulations to show that many of the currently used approaches are inappropriate for accurately estimating the intrinsic growth rate *µ*. We quantify the errors for the intrinsic growth rate *µ* when the maximum attained growth rate *µ*_max_ is used as an estimator and the generating model is known. Next, we present the limited set of conditions in which an exponential approximation can be used for estimating *µ*. Importantly, we demonstrate that using inaccurate estimates of *µ* to estimate the relative fitness often leads to inaccurate fitness estimates and sometimes to wrongly classifying a beneficial mutation as deleterious in some cases. Finally, we apply our theoretical insights to previously published data and show that both absolute and relative growth rate estimates may vary greatly depending on the method. Overall, we present a systematic evaluation of different methods, with recommendations for best practices.

## 2 Results & Discussion

### 2.1 Literature review: How does the community analyze the data?

We reviewed 50 papers from evolution and ecology that estimated growth curves for all types of microbial data (see Table S1 and Methods section). Most of the data (90%) were acquired by an automated microplate reader tracking optical density (OD) over time. Other data types included cell counts or fluorescent yields over time. Several papers (6%) did not report the starting inoculum size. Of those papers that reported the inoculum size, 52% used a fixed absolute initial population size 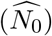 for inoculation whereas 44% used a constant initial population fraction 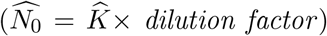, which we hereafter refer to as the dilution fraction. A constant dilution fraction means that stationary-phase cultures, whose populations are at their carrying capacity, *K*, were diluted by a constant dilution factor (e.g., 1/1000×). This means that for experiments with a constant dilution fraction the absolute population size of the inoculum, *N*_0_, differed between strains/treatments when the carrying capacities, *K*, were different. For experiments using a constant dilution fraction, the reported dilution factor varied between 10^−4^ to 10^−1^ with a geometric mean value of 10^−2.37^.

We found that the growth rate was by far the most commonly estimated growth parameter (94% of all papers reviewed). The other estimated growth parameters were: the carrying capacity (44%), the lag time (34%), and the area under the curve (AUC; 12%). The growth rate was usually reported as an absolute value for each strain/treatment. Moreover, 24% of papers estimated the relative growth rate, or relative fitness, of different strains as compared to the wild-type.

Methods that explicitly fit a model of population growth were used about as often as “model-free” or “nonparametric” approaches (i.e., methods that do not require a model; Figure 2A). One particular model-free approach, the “Easy Linear” method, was especially popular (right doughnut chart of Figure 2A): it was used in about a third of all papers. When models *were* used, only one model was usually reported to have been fit (left doughnut chart of Figure 2A).

**Figure 2:**
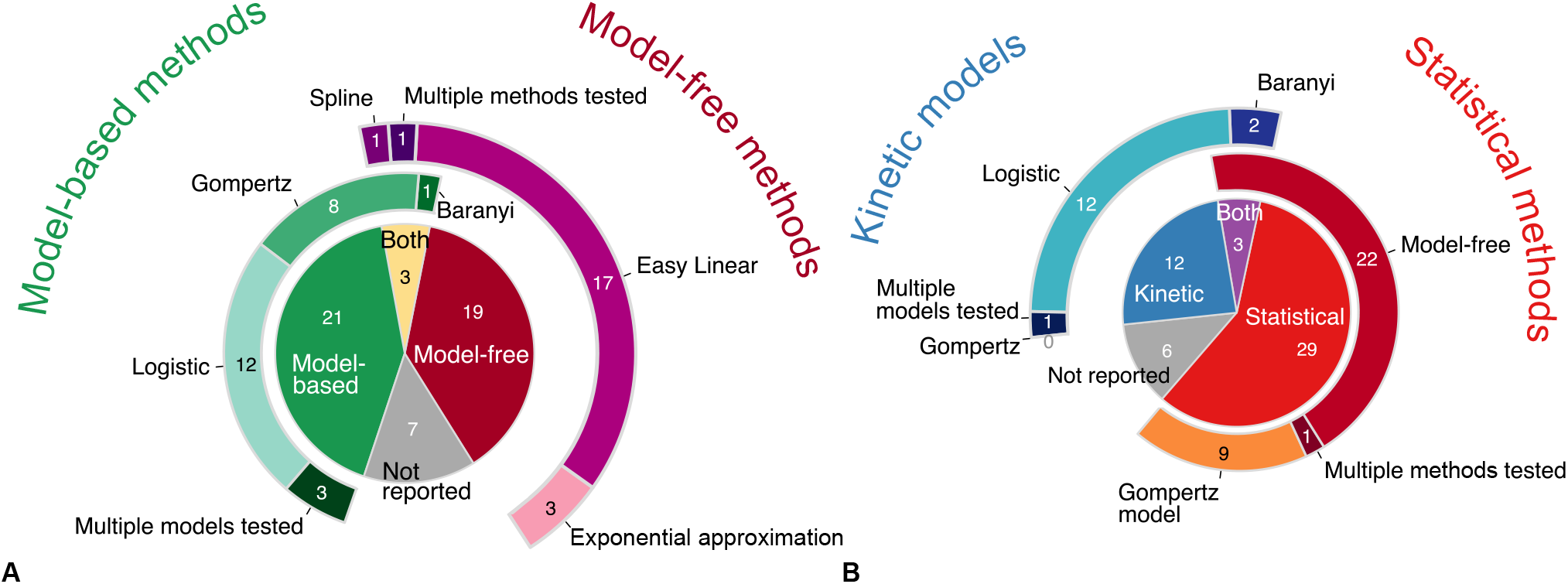
Pie and doughnut charts of literature review results. **A) Model-free methods were used about as often as model-based methods** across all 50 papers reviewed. “Both” refers to when both a model-free and a model-based method were used in the same paper. The doughnut charts surrounding the central pie-chart illustrate how frequently different models, including both statistical and kinetic models, were used (left, shades of green) and how frequently different statistical model-free methods were used (right, shades of purple). **B) Statistical approaches were used more often than kinetic models**. “Both” refers to when both a kinetic model(s) and a statistical approach(es) were used in the same paper. The doughnut charts surrounding the central pie-chart illustrate how frequently different kinetic models were used (left, shades of blue) and how frequently different statistical approaches were used (right, shades of red-orange). The Gompertz model is listed twice because it is either a kinetic model or a statistical model, depending on the equation; however, no paper was found to use a kinetic Gompertz model. Both of the statistical Gompertz models listed in Table 1, Gompertz and modified Gompertz, are grouped together. For more details on the methods and equations used by each paper, see table S1. All slices within each pie and doughnut chart show the counts of papers included.

We classified the growth curve analysis methods as either kinetic or statistical (Figure 2B). A kinetic model is a mechanistic representation that allows researchers to simulate the underlying process. In contrast, a statistical approach enables researchers to describe and quantify the pattern of interest but without simulating the underlying process. We found that statistical approaches were used more often than kinetic models (Figure 2B). The most popular methods within the statistical approach were the various model-free methods (right doughnut of Figure 2B). The logistic model was by far the most popular kinetic model used (left doughnut of Figure 2B). Depending on its equation, the Gompertz model is either a statistical or a kinetic model (table 1); however, we found that the kinetic Gompertz model was never fitted whereas the statistical Gompertz model was popular (18% of all papers and 28% of all statistical methods used).

**Table 1:** Different population growth models: The population growth models considered in this paper with their equation and parameters. Kinetic models are indicated by (Kin) and describe the population size *N* as a function of time *t*. Every kinetic model includes a carrying capacity *K*, an initial population size *N*_0_ and an intrinsic growth rate *µ*. The kinetic Richards model includes a parameter *β* to adjust its inflection point and the Baranyi model has a lag phase defined by *h*_0_. The Huang model has both with parameters *α* and *τ*, respectively. Statistical models are indicated by (S) and describe *y* = ln(*N/N*_0_) as a function of time *t*. Every statistical model includes a carrying capacity *A* = ln(*K/N*_0_), a maximum growth rate *µ*_max_ and a lag phase *λ*. The modified Gompertz model (S) displays a second increase after the growth reaches a first saturation plateau. The parameters *t*_*shift*_ and *α* control the time and the slope of the second increase. The inflection point in the statistical Richards model is adjustable by the parameter *v*.

| Model | Equation | Parameters |
| --- | --- | --- |
| Logistic (Kin) | $N(t) = \frac{N_0 K \exp(\mu t)}{K + N_0 (\exp(\mu t) - 1)}$ | $K, N_0, \mu$ |
| Gompertz (Kin) | $N(t) = (N_0/K)^{\exp(-\mu t)} K$ | $K, N_0, \mu$ |
| Richards (Kin) | $N(t) = \frac{N_0 K}{(N_0^\beta + (K^\beta - N_0^\beta) \exp(-\mu \beta t))^{-1/\beta}}$ | $K, N_0, \mu, \beta$ |
| Baranyi (Kin) | $N(t) = \frac{(-1 + \exp(h_0) + \exp(\mu t)) N_0 K}{(\exp(\mu t) - 1) N_0 + \exp(h_0) K}$ | $K, N_0, \mu, h_0$ |
| Huang (Kin) | $N(t) = \frac{N_0 K}{N_0 + (K - N_0)(1 + \exp(\alpha \tau))^{\mu/\alpha} (\exp(\alpha t) + \exp(\alpha \tau))^{-\mu/\alpha}}$ | $K, N_0, \mu, \alpha, \tau$ |
| Logistic (S) | $y(t) = \frac{A}{1 + \exp(4\mu_{\max}(\lambda - t)/A + 2)}$ | $A, \mu_{\max}, \lambda$ |
| Gompertz (S) | $y(t) = A \exp(-\exp(\mu_{\max} \exp(1)(\lambda - t)/A + 1))$ | $A, \mu_{\max}, \lambda$ |
| modified Gompertz (S) | $y(t) = A \exp(-\exp(\mu_{\max} \exp(1)(\lambda - t)/A + 1)) + A \exp(\alpha(t - t_{shift}))$ | $A, \mu_{\max}, \lambda, \alpha, t_{shift}$ |
| Richards (S) | $y(t) = A(1 + \nu \exp(1 + \nu + \mu_{\max}(1 + \nu)^{1+1/\nu}(\lambda - t)/A))^{-1/\nu}$ | $A, \mu_{\max}, \lambda, \nu$ |

We found that many growth curve experimental methods, data, and analyses do not yet conform with recommendations for reproducible research (e.g., Wilkinson et al. 2016; MunafÒ et al. 2017). Over 10% of the papers reviewed reported insufficient information about how growth curves were analyzed and therefore could not be classified for Figure 2. Some highlycited papers (e.g. Gullberg et al. 2011; Trindade et al. 2012) neither cited an established method nor included sufficient description of their *ad hoc* methods for estimating growth rates (see table S1). Beyond reporting of experimental methods, the data-set itself was often not shared: about half (46%) of all papers do not show any figures of nor provide any of the growth curve data (see table S1). 40% of papers provided plots of at least a subset of the growth curves and 14% of all papers published their raw growth curve data.

Our finding from Figure 2 that ∼ 13% of articles provide insufficient information regarding their growth curve analysis methods likely *under*estimates the magnitude of the problem. This is because we found articles for inclusion in the review by searching among the citations to previously published growth curve analysis methods papers (see methods). Therefore, most of the papers we included cited an established method for analyzing growth curve data. Hopefully these issues of methods under-reporting will improve as scientists become more knowledgeable about recommendations for open science and data management (Wilkinson et al., 2016; Munafò et al., 2017).

Our finding of insufficiently reported information regarding the analysis of growth curves corroborates previous concerns about the lack of a standard method for growth curve analysis (Fernandez-Ricaud et al., 2016). The remainder of our article discusses different methods for analyzing growth curves and, thus, will hopefully contribute to an increased appreciation of why it is important to provide sufficiently detailed methodological information on data analyses.

### 2.2 Exposition of existing models and methods

#### 2.2.1 Conceptual distinctions between parametric vs non-parametric methods and statistical vs kinetic models

A variety of approaches have been developed over the years to describe and quantify growth curves, as shown in Figure 2. Below we explain the main differences between the most commonly used approaches and models, then we compare the advantages of each.

##### Model-free vs model-based methods

One way to classify the different methods is to distinguish between model-free (or non-parametric) methods and model-based (or parametric) methods. Model-free methods use an algorithm to find an estimate of the growth rate that is relatively robust to any noise error in the data. For example, in the classical exponential approximation from Monod (1949) that is featured in many introductory microbiology textbooks, growth rate is estimated by measuring cell concentrations (*N*_1_ and *N*_2_) at two time points (*t*_1_ and *t*_2_) during the “log” phase of growth, then calculating *R* = (log_2_ *N*_2_ − log_2_ *N*_1_)*/*(*t*_2_ − *t*_1_). This is an algorithm that can easily be used by hand or performed by a computer. The Easy Linear method (Hall et al., 2014; Mira et al., 2017) is a more complex algorithm that uses a sliding window of five successive data points to calculate the maximum slope among many linear regressions fitted to the log-scale growth curve data. Another example is the Spline method that calculates the maximum value of the first derivative of the log-scale growth curve data by either using the mean of three successive pairs of points (e.g., Ashino et al. 2019) or kernel smoothing of the growth curve (e.g., Kahm et al. 2010; Petzoldt 2020) in order to remove experimental noise. The parameters of these algorithms are usually tunable, for example the size of the sliding window used by the Easy Linear method. However, no explicit assumptions are made about the shape of the growth curves.

Model-based methods, on the other hand, use equations to explicitly describe the relationship between time, the independent variable, and population size (or a proxy of population size, like OD), the dependent variable. A model fitting algorithm is then used to find the model parameters that best fit the observed data, usually by numerical minimization of the residual sum of squares. Model-based methods tend to be preferred by theoretical and statistical biologists because models specifically define the assumptions that are being made, model-based methods have defined protocols for assessing goodness of fit, and model-fitting allows quantification of the (frequentist or Bayesian) error of the estimates (Otto and Day, 2011; Bolker, 2008).

Empiricists tend to prefer to apply model-free approaches to biological growth curve data both because it is technically more difficult to fit and compare models and because of the inflexibility of existing models to fit the data (source: personal communication). In other words, the main advantage of model-free approaches is that they do not require a model. Model-free approaches have drawbacks, however: since there is no model, it is not clear how to compare the likelihood or goodness-of-fit between different methods and bootstrappind is necessary to quantify the error around estimates (for example, in order to generate the 95% confidence intervals).

##### Kinetic vs statistical models

Within model-based methods, there is a distinction between kinetic and statistical models. Kinetic models may also be called mechanistic/process models and statistical models can be called phenomenological/pattern models (Bolker, 2008). We define kinetic models as mechanistic representations that simulate the underlying process of interest. For population growth curve models in particular, the *per capita* growth rate of a kinetic model is by definition the intrinsic growth rate *µ* when the cell’s resources are limitless or otherwise optimal.

Statistical models can be thought of as ‘black-box empirical models’ (Chezeau and Vial, 2019). They are created by selecting functions that have a similar shape as the pattern of interest. Then, the parameters of those functions are given a biologically relevant meaning. For growth curves in particular, statistical population growth models are defined such that the point on the curve with the fastest rate of *per capita* growth (i.e., the inflection point of *y* = ln(*N/N*_0_)) corresponds to the maximum growth rate *µ*_max_ (Zwietering et al., 1990, equations 4 & 5). Model-free (non-parameteric) methods all belong to the statistical category, as shown in Figure 2B, as they quantify specific parameters of interest without simulating the underlying process.

Although the distinction between kinetic models and statistical methods may seem arcane, we explain below how serious pitfalls in estimating growth rates are a direct result of the differences between growth rates estimated by kinetic models (i.e., the intrinsic growth rate *µ*) and statistical methods (i.e., the maximum growth rate *µ*_max_).

#### 2.2.2 Mathematical description of models and model parameters, including the initial fraction

Many models have been developed to describe growth curves (e.g., Tsoularis and Wallace 2002; Huang 2011; Baranyi and Roberts 1994). We have summarized the equations and parameters of the most prevalent models found by our literature review in Table 1, distinguishing kinetic models (Kin) from statistical models (S). As explained in the section above, a main difference to note is that kinetic models are defined in terms of the intrinsic growth rate *µ*, whereas statistical models are defined in terms of the maximum growth rate *µ*_max_. Kinetic models describe population size *N* as a function of time (i.e., linear scale), whereas statistical models describe *y* = ln(*N/N*_0_) as a function of time (i.e., they operate on a logarithmic scale).

The common feature of all models, whether kinetic or statistical, is that they have a sigmoid or ‘S’ shape (Zwietering et al., 1990). All models consider single-strain, well-mixed bacterial populations whose every individual divides at the same *per capita* rate, although the division rate varies over time. Each of these populations starts with *N*_0_ microbes and their maximum population size is defined by a carrying capacity, written as *K* for kinetic models or *A* for statistical models.

The models differ in important ways. Most of the models derive from the logistic model but seek to generalize it (Tsoularis and Wallace, 2002). For example, the Richards (both kinetic and statistical) and Baranyi models include a parameter to set the growth inflection and a lag phase, respectively, whereas the Huang model incorporates both. The Gompertz models (both kinetic and statistical) have no additional parameters but display faster growth than the logistic model for the same set of parameters since they have a higher *per capita* growth rate. Futhermore, the Richards model requires an extra parameter *β* that determines how quickly deceleration occurs as the stationary phase is reached, whereas the Baranyi model includes a lag phase specified by the parameter *h*_0_. The Huang model includes a lag phase defined by *τ* in addition to a parameter *α* determining the curvature.

The initial fraction, *N*_0_*/K*, is an important value to keep in mind throughout our paper. This is the size of the population at inoculation (i.e., *N* at time *t* = 0) divided by the final carrying capacity. Only when the initial fraction is small can one distinguish models that have an initial exponential growth, like the logistic model, from models with a lag phase, like the Gompertz, Richards, Baranyi, and Huang models.

#### 2.2.3 What is the difference between *µ* and *µ*_max_?

It is important to distinguish between three growth rate estimators: the maximum population growth rate (max (d*N/*d*t*)), the *per capita* maximum growth rate (*µ*_ma x_), and the *per capita* intrinsic growth rate (*µ*). The maximum population growth rate max (d*N/*d*t*) is the fastest increase in size achieved by the *entire population*. We are not interested in the maximum population growth rate parameter since it is not estimated by any of the methods or models we discuss here; we only mention it so that the reader does not mistake it for the maximum growth rate (*µ*_max_). The maximum growth rate *µ*_max_ is the fastest *per capita* number of divisions per unit of time actually achieved in the observed growth curve. In more quantitative terms, *µ*_max_ is the maximum value of the curve d ln(*N/N*_0_)*/*d*t* (Figures 1B-C). As mentioned above, the maximum growth rate *µ*_max_ is a value estimated using statistical approaches. Finally, the intrinsic growth rate *µ* (sometimes denoted as *r* (Sprouffske and Wagner, 2016), called the Malthusian parameter of population growth, or the intrinsic rate of increase) is the fastest *per capita* number of divisions per unit of time theoretically possible and, because it is a kinetic model parameter, it is used for simulating population growth processes. We here focus on the intrinsic growth rate *µ* and the maximum growth rate *µ*_max_ because these are the two quantities estimated by the most used methods.

There is an important conceptual difference between the intrinsic growth rate *µ* and the maximum growth rate *µ*_max_. The intrinsic growth rate *µ* is the theoretical maximum number of cell divisions per time unit assuming population dynamics that follow an exponential law. However, No real population achieves an infinite size because the division process is limited by space and/or nutrients, for instance. Thus, the number of divisions per time unit is not constant over time, so the maximum division rate *µ*_max_ is the largest *per capita* value observed during the population growth. Therefore, the intrinsic growth rate *µ* can quantify the strain-specific division rate theoretically independently of the environment or experimental conditions. On the other hand, the maximum growth rate *µ*_max_ (like other values estimated by statistical methods) is always specific to the experiment itself and cannot be generalized as a strain-specific value that applies to different environments or conditions. As will be shown below, in the best case scenario *µ*_max_ approximates *µ*, but in other scenarios *µ*_max_ is a composite parameter that depends on other values, like the inoculum size and lag time.

Previous work has pointed out confusions between different growth rate estimators (Perni et al., 2005). The confusion between these terms is so prevalent that some papers mistook *µ* for *µ*_max_ (Yang et al., 2006), vice versa (Wu et al., 2017), or distinguished between the two but swapped the names (Khan et al., 2017). Furthermore, some authors wrote the kinetic logistic equation as d*N/*d*t* = *µ*_max_(1 − *N*_0_*/K*)*N* (Petzoldt, 2020), whereas other authors preferred d*N/*d*t* = *µ*(1 − *N/K*)*N* and *µ*_max_ = max(d ln(*N* (*t*)*/N*_0_)*/*d*t*) (Sprouffske and Wagner, 2016). Different naming conventions become even more misleading for models in which the intrinsic growth rate is a function of the resource concentration, such as the Monod class of models that have their own specific, kinetic definition for *µ*_*max*_ (Monod, 1949; Chezeau and Vial, 2019). We will not discuss substrate-use models herein. Having explained the conceptual differences between the *µ*_max_ maximum growth and *µ* intrinsic growth rates above, we now expand on the mathematical differences.

##### Deriving the difference between *µ*_max_ vs. *µ*

In the following, we mathematically explain why *µ*_max_ is not always a good proxy for *µ*, especially at large initial population fractions, *N*_0_*/K*. Analytical math is combined with simulations to show for which initial population fractions an experimenter can estimate the intrinsic growth rate *µ* from the maximum growth rate *µ*_max_.

We assumed that the population dynamics follow one of the kinetic growth models from Table 1 and mathematically derived *µ*_max_ for the five kinetic models. As reported in Table 2, *µ*_max_ depends on the system parameters, namely the initial population fraction *N*_0_*/K* as well as the parameters *β* and *h*_0_ for the Richards and the Baranyi models, respectively.

**Table 2:** Maximum growth rates: The maximum growth rate *µ*_max_ for the population growth models considered in this paper. Kinetic models are indicated by (Kin), whereas the statistical models are indicated by (S). The maximum growth rate was derived for the kinetic models by analytically determining max(d*y/*d*t*) = max(d ln(*N/N*_0_)*/*d*t*). All statistical models have the same maximum growth rate, whereas the maximum growth rate differs between kinetic models.

| Model | Derived maximum growth rate $\mu_{\max}$ |
| --- | --- |
| Logistic (Kin) | $\mu(1 - N_0/K)$ |
| Gompertz (Kin) | $-\mu \ln(N_0/K)$ |
| Richards (Kin) | $\mu(1 - (N_0/K)^\beta)$ |
| Baranyi (Kin) | $\frac{\mu e^{-h_0}(-2(1 + \sqrt{(-1 + e^{h_0})(e^{h_0}(K/N_0) - 1)})(N_0/K) + e^{h_0}(1 + N_0/K))}{1 - N_0/K}$ |
| Huang (Kin) | Numerical solution |
| Logistic (S) | $\mu_{\max}$ |
| Gompertz (S) | $\mu_{\max}$ |
| modified Gompertz (S) | $\mu_{\max}$ |
| Richards (S) | $\mu_{\max}$ |

The results of Table 2 are illustrated by the points in Figure 3A. The estimated maximum growth rate (*µ*_max_) values differ between models with the same parameter values (intrinsic growth rate *µ* = 1 and carrying capacity *K* = 10^5^). In general, the maximum growth rate is approximately equal to the intrinsic growth rate when the initial fraction of individuals satisfies *N*_0_*/K* « 1 and (*N*_0_*/K*)^*β*^ « 1 in the Logistic and Richards models, respectively. Indeed, these conditions lead to *µ*_max_ = *µ*(1 − *N*_0_*/K*) ≈ *µ* and *µ*_max_ = *µ*(1 − (*N*_0_*/K*)^*β*^) ≈ *µ* for the Logistic and Richards models, respectively (see Table 2). The Gompertz model is a special case since the maximum division rate is a good proxy for the intrinsic division rate when the initial fraction is large, roughly equal to exp(−1) ≈ 0.37 (i.e., an inoculum size corresponding to a dilution factor for the stationary phase batch culture of between one-third and two-fifths).

**Figure 3:**
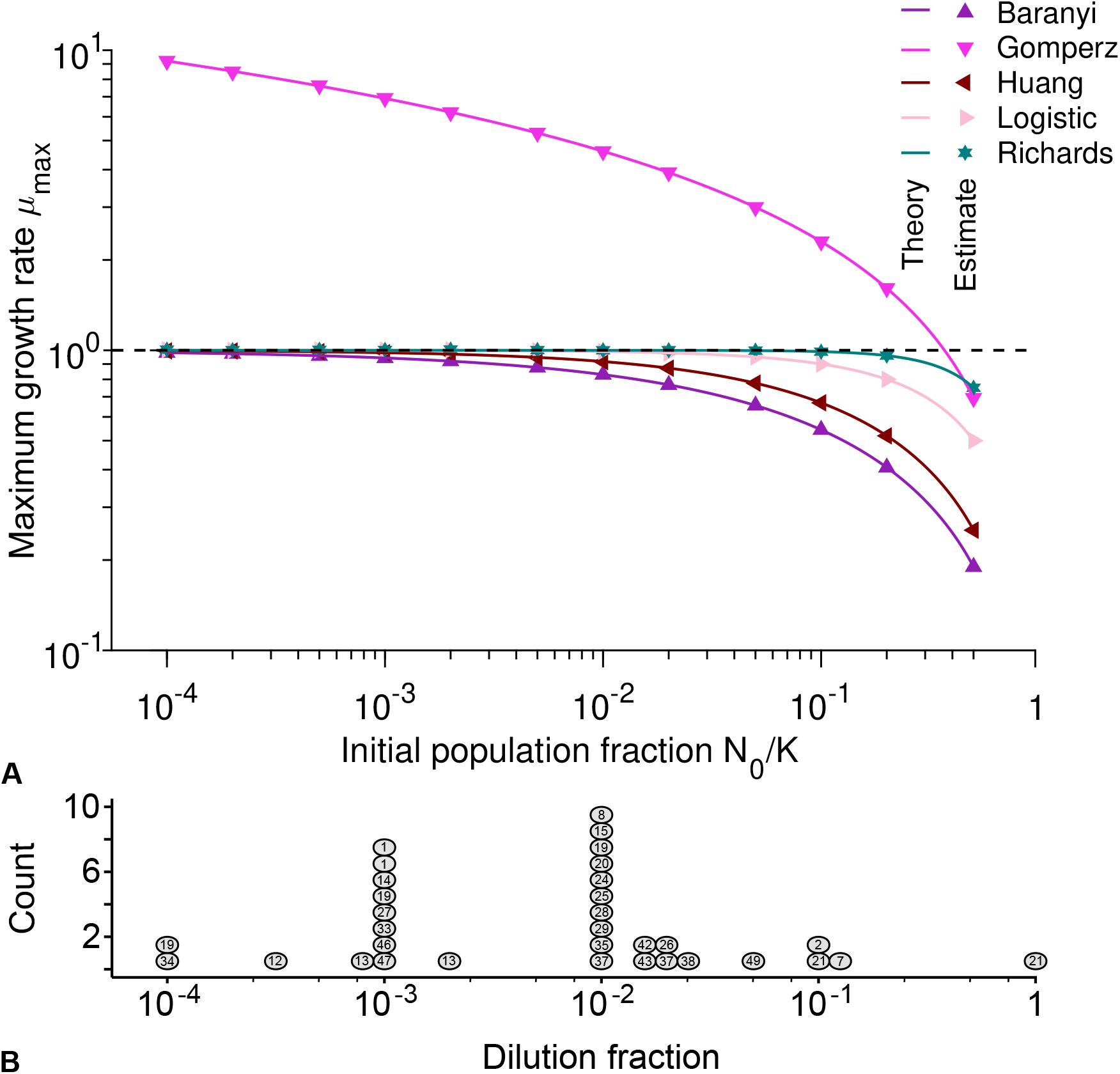
**A) The Gompertz model and large initial population fractions make the maximum growth rate a poor proxy for the intrinsic growth rate:** Maximum growth rate *µ*_max_ versus initial population fraction *N*_0_*/K* for different kinetic population growth models, where *µ* = 1. Each point represents estimated values by Spline (from Petzoldt 2020) from simulated data averaged over 10^4^ stochastic realizations. The solid lines correspond to the analytical predictions of the maximum growth rate (see Table 2). The dashed line shows the intrinsic growth rate value *µ*. Parameter values: *K* = 10^5^, *µ* = 1, *α* = 2, *β* = 2, *h*_0_ = 2 and *τ* = 2. The initial population fraction is defined as a combination of model parameters, *N*_0_*/K*, while the initial dilution fraction is an empirical quantity extracted from experimental data as detailed in the literature review methods. **B) For the most commonly used dilution fractions in the literature, the maximum growth rate is a good proxy for the intrinsic growth rate:** Histogram of the estimated dilution fractions observed for the 27 (out of 50) papers that provided sufficient information to estimate this value. Each circle represents a publication and the number inside the circle indicates the number of the reference as given in supplementary table S1. Several publications appear more than once because they used more than one dilution fraction for different experiments.

In order to test the analytical predictions from Table 2, we evaluated the growth rates as estimated by model-free methods using data simulated under each of the five population growth models. Unlike experimental data, for which the true *µ* value that generated the data is never known, estimating the growth rate from simulated data allows us to check the accuracy of the estimates as compared to the known *µ* parameter that the data was simulated under.

We focus on two model-free methods, the popular Easy Linear (Hall et al., 2014) and the Spline (e.g., Adkar et al. 2017; Ashino et al. 2019) methods, to determine the maximum growth rate *µ*_max_. Both methods assume that only the exponential stage of growth is useful to estimate the maximum growth rate. We generated data using individual based stochastic simulations for the Gompertz, Richards, Logistic, Huang, and Baranyi models. Then we used the two different model-free methods, Spline and Easy Linear, to compute the maximum growth rate *µ*_max_ for different parameter values. In practice, both model-free methods provided us with the same results. Our simulated data averaged over several stochastic realizations did not include the myriad sources of noise present in experiments, therefore resulting in a low noise level.

As shown by the lines in Figure 3A, there is an excellent agreement between our analytical predictions (lines) and the estimates from simulated data (points). As predicted analytically, the estimated maximum growth rate *µ*_max_ is not equal to the known intrinsic growth rate *µ* value used to create the simulations, unless *N*_0_*/K* « 1. For the Baranyi, Huang, Logistic, and Richards models, the smaller the initial population fraction, the better the maximum growth rate performs as a proxy for estimating the intrinsic bacterial growth rate, *µ*. However, this is not the case for the Gompertz model. For the Baranyi and Richards models (supplementary Figures S4A and S4B), the smaller the parameter *h*_0_ and the larger the parameter *β*, the closer is the maximum growth rate *µ*_max_ to the intrinsic growth rate *µ*. Similarly, for the Huang model, the higher the curvature defined by *α* and the shorter the duration *τ* of the lag phase, the better *µ*_max_ is as a proxy for *µ* (see Figures S4C and S4D).

We have shown that the maximum growth rate *µ*_max_ is not always equivalent to the intrinsic growth rate *µ*. Therefore, methods that estimate the maximum growth rate *µ*_max_ but then (often implicitly) assume that this value can be treated as the *µ* of a kinetic model must be applied with caution. As we have demonstrated in Table 2 and Figure 3A, *µ*_max_ tends to *under*estimate the true intrinsic growth rate *µ* – except when population growth follows the Gompertz kinetic model, in which case the maximum growth rate *µ*_max_ mostly *over*estimate the true intrinsic growth rate *µ*. This is because the *per capita* growth rate is generally smaller than the intrinsic one. Hence, we recommend that a clear distinction must be made between the intrinsic growth rate *µ* and the maximum growth rate *µ*_max_.

##### The initial fraction is a key parameter determining the relationship of µ_max_ and µ

Above we showed that (for most models) we can use the estimated maximum growth rate *µ*_max_ as an approximation of *µ* when initial population fractions *N*_0_*/K* are small. We estimated the initial dilution fractions used in different experiments in order to ascertain whether most studies are using appropriately small initial fractions. Figure 3B shows the estimated dilution fractions used by different papers included in our literature review. Papers that had experiments with multiple, different batch culture starting conditions are included as multiple points with the same number. For example, Ganucci et al. 2018 (labeled as 21) has a dilution fraction near 1. The authors used cell viability counts to track yeast growth in media with increasing ethanol concentrations, sometimes resulting in almost no growth. Two values from Ganucci et al. (2018) are summarized in Figure 3B, indicating the largest (0.1) and the smallest (0.94) dilution fractions observed. Since methods differ between publications, with some using a mid-exponential phase culture and others using a stationary phase culture for inoculation (see supplementary table S1), the estimated dilution fraction should be considered as an upper bound for the initial fraction used in each paper.

More than two-thirds of papers use at least one estimated dilution fraction smaller than 10^−2^; the geometric-mean observed dilution fraction was 10^−2.7^. For such small dilution fractions, if there is no lag time, and if growth follows one of the population growth models except Gompertz, the maximum growth rate *µ*_max_ tends to be a good estimator of the intrinsic growth rate *µ*. When the true growth curve dynamics in the experiments follows one of the kinetic models other than Gompertz, the relative difference between the maximum growth rate and the intrinsic growth rate for the mean initial population fraction *N*_0_*/K* = 10^−2.7^ is between 0-12%. Here, the largest difference between *µ*_max_ and *µ* is obtained for the Baranyi model. However, for the Gompertz model the relative difference between the maximum growth rate and the intrinsic growth ranges from -821% to 31% (see Figure 3).

##### Discussion of the difference between *µ*_max_ and *µ*

We showed that the *µ*_max_ values calculated from kinetic models depend on other parameters in addition to *µ*, such as the initial population fraction. Conversely, one cannot obtain *µ* from *µ*_max_ alone. For kinetic models, additional parameters such as the initial population size and carrying capacity are required (see Table 1) to be able to calculate *µ* from *µ*_max_. For statistical approaches, which are specified directly in terms of *µ*_max_, *µ* is not defined. Nevertheless, even when *µ*_max_ is estimated by statistical models, its estimated value will be different when experimental quantities such as the initial population size and carrying capacity change. In reality, the estimated values for both *µ*_max_ and *µ* may also vary with the experimental conditions (such as genotype, medium, temperature, etc).

We emphasize that the main difference between the maximum growth rate *µ*_max_ and the intrinsic growth rate *µ* is that *µ*_max_ is a statistical quantity, whereas *µ* is a model parameter. Therefore, obtaining different estimates of *µ* and *µ*_max_ is expected and not a sign of bad performance of a model or method, especially for large initial population fractions. For example, the manual of one software (Delaney, 2014) provides options for fitting various statistical models and a single kinetic model but discourages users from applying the kinetic model because “it predicts fastest growth” as compared to the other implemented models. Our results can readily explain this observation and debunk the implied worse performance of the kinetic model. Indeed, the publication associated with this software recommends a large inoculum size (Delaney et al. 2013; associated studies labeled as 8, 22, 25, 42, and 43 in Figure 3B and supplementary table S1), for which we showed that *µ*_max_ consistently overestimates *µ*.

Given the differences between *µ*_max_ and *µ*, which of these should preferably be estimated? Unfortunately, there is not a one-size-fits-all answer to this question. From a theoretician’s perspective, we recommend that researchers estimate the intrinsic growth rate *µ*. Most studies perform growth curve experiments to characterize growth for specific strains, treatments, or environments in the (either explicit or implicit) context of population ecology or mechanistic models – all of which are defined in terms of *µ*. Only the intrinsic growth rate *µ* can be used for simulating mechanistic models and therefore to identify the mechanisms which best explain the observed dynamics. The maximum growth rate *µ*_max_, on the other hand, is phenomenological: it describes what is observed in the data contingent on the population starting conditions and the experimental environment. At best, *µ*_max_ approximates *µ*. However, from an experimenter’s perspective, estimating *µ*_max_ has the advantage that model-free methods are technically easier to use than estimators of kinetic models, especially for data that display diphasic or other non-sigmoid/non-’S’ shaped growth. We strongly urge researchers who decide to estimate *µ*_max_ to use (and report) a small initial population fraction and to assert that the data do not have a significant lag time.

When using statistical methods, a second decision is necessary about whether to use model-based or model-free methods. In our noise-free, simulated data, model-based and model-free statistical methods yielded the same estimates for *µ*_max_ (Figure 3A); future work should elaborate on their performance in the presence of different sources of noise (e.g., to expand on the work that has already been done by Mira et al. (2017) for the Easy Linear method). Although models are preferable from a theoretician’s view, certain types of experimental data (for example, displaying diauxic growth, curves without samples in the stationary phase, and other non-sigmoid shaped data as well as very noisy data) may not be fitted well by a model of sigmoidal growth. In this case the data may be better summarized by a model-free statistical method. Importantly, we recommend that the choice of method is justified clearly and in writing, no matter which method is used.

### 2.3 Guidelines for estimating growth rates

#### 2.3.1 Ad hoc fitting an exponential curve to growth curves should be avoided

One common approximation (Monod, 1949; Kassen, 2014) that is used to obtain an estimate of the intrinsic growth rate *µ* is to fit an exponential equation, like 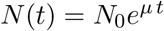, to the early phases of growth (i.e., during the “log”/”linear” phase, prior to the deceleration and stationary phases). This method is explained in many introductory microbiology textbooks, as previously summarized in the explanation of model-free methods of section 2.2.1 above, and we refer to this approach as an exponential approximation. Under the exponential approximation, the intrinsic growth rate is given by *µ* = ln(*N* (*t*)*/N*_0_)*/t*.

To test the accuracy of the exponential approximation when applied to batch culture population growth, we expressed the intrinsic growth rate as a function of population size and other possible parameters for different kinetic population growth models (Table 3 and Figure S5).

**Table 3:**
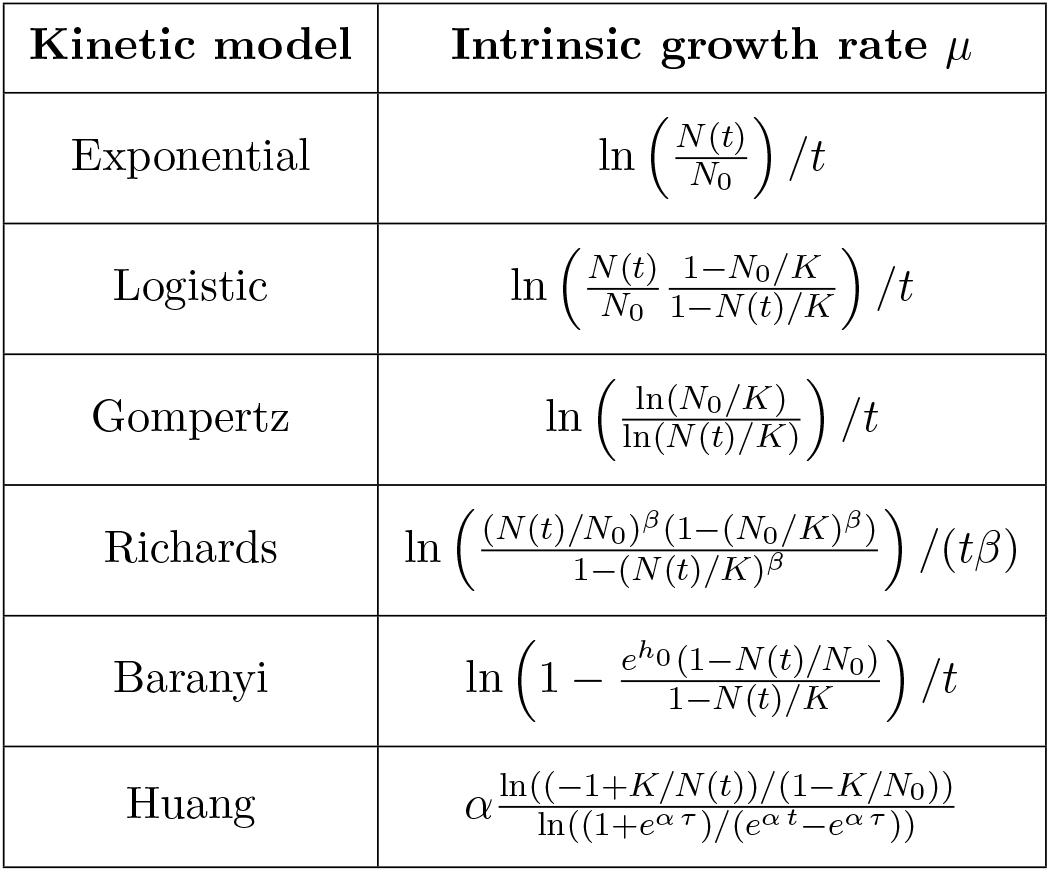
Intrinsic growth rate as function of the model parameters at time *t* for different kinetic population growth models. The difference between the equations indicates that the exponential approximation may often not be appropriate; see also Figure S5.

We found that the exponential approximation is frequently a poor estimator of the intrinsic growth rate *µ*. The exponential approach is never valid for a population following Gompertz growth. There is no parameter range for which the equation ln (ln(*N*_0_*/K*)*/* ln(*N* (*t*)*/K*)) */t* reduces to ln (*N* (*t*)*/N*_0_) */t* (see Table 3). The exponential approach is valid for the logistic growth when the initial population size is very small in comparison to the carrying capacity (i.e., *N*_0_ « *K*) and for time points at which the population size remains small in comparison to the carrying capacity (i.e., *N* (*t*) « *K*). These conditions lead to ln(*N* (*t*)(1 − *N*_0_*/K*)*/*(*N*_0_(1 − *N* (*t*)*/K*)))*/t* ≈ ln(*N* (*t*)*/N*_0_)*/t* (see Table 3). This makes sense since the phase during which these conditions are satisfied corresponds to the regime in which logistic growth can be reduced to exponential growth. The same conditions apply to Baranyi growth, with the additional condition that the lag phase must be short (i.e., *h*_0_ « 1), so that one obtains ln 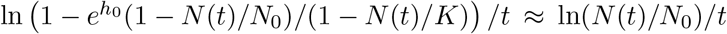 (see Table 3). If the lag phase is not short, the exponential phase starts later whereas the exponential approach assumes that it starts at the beginning of the growth. Richards growth is more complex. Here, the quantities (*N*_0_*/K*)^*β*^ and (*N* (*t*)*/K*)^*β*^ must be much less than 1 to make the exponential approach valid. Thus, the larger the deceleration parameter *β* is when *N*_0_ « *K* and *N* « *K*, the more abruptly the “log”/”linear” phase transitions into the stationary phase, and the more valid the exponential approximation becomes, so that ln (*N* (*t*)*/N*_0_)^*β*^(1 − (*N*_0_*/K*)^*β*^)*/*(1 − (*N* (*t*)*/K*)^*β*^) */*(*tβ*) ≈ ln(*N* (*t*)*/N*_0_)*/t* (see Table 3). Consequently, the exponential approximation is valid only in a very restricted set of conditions: when there is no lag phase, the initial population fraction is very small, and the measured population sizes remain small as compared to the carrying capacity. Most experimental data probably do not meet this necessary set of conditions.

It is of note that throughout the literature, including introductory textbooks, the term “exponential growth rate” tends to be used to describe the intrinsic growth rate *µ* and sometimes deemed the same as the maximum growth rate *µ*_max_ (e.g., Basra et al. 2018; Novak et al. 2009). We here strongly caution against this conflation of potentially very different quantities. In particular, we recommend that approximating batch culture growth with an exponential curve in order to get an estimate for the intrinsic growth rate *µ* requires careful assurance that the assumption is valid for the data at hand.

#### 2.3.2 Theory predicts that using *µ*_max_ or the exponential approximation for estimating relative fitness can yield wrong results

In evolution and population biology, the relative fitness *w* of a mutant (M) compared to a wild-type (WT) is classically defined as the ratio of *µ*^*M*^ */µ*^*WT*^. The relative fitness or the selection coefficient, defined as *s* ≡ *w* − 1 = *µ*^*M*^ */µ*^*WT*^ − 1, is used to classify a mutant as deleterious (*w <* 1 or *s <* 0), neutral (*w* = 1 or *s* = 0), or beneficial (*w >* 1 or *s >* 0). Thus, to infer how natural selection favors one strain over another from monoculture growth curves, microbial ecologists and evolutionary biologists need the intrinsic growth rate, which is obtained by fitting a kinetic model.

One argument regarding the issues of miscalculation of the intrinsic growth rate *µ* discussed above is that these concerns are important for *absolute* growth rate estimates, but can be disregarded when considering *relative* estimates. Here, the argument is that whereas absolute estimates cannot be compared between data-sets, relative growth rates can be estimated within a data-set by using a common reference sample and these relative growth rates can then be compared between data-sets.Moreover, with regards to selection coefficients, the sign may be more important than the absolute value. Namely, incorrect absolute estimates should yield correct rankings of growth rate estimates, and thus correctly estimated signs of the selection coefficient. Below, we demonstrate that these assumptions are wrong and that incorrect estimates of the growth rates (for example, by assuming that *µ*_max_ = *µ*) can severely affect the classification of strains into beneficial, neutral, or deleterious.

To demonstrate the effects on relative fitness estimates of assuming *µ*_max_ = *µ*, we simulated growth curve data from separate batch monocultures for two strains and estimated their maximum growth rates *µ*_max_ using the growth curves. Then we assumed (erronously) that the maximum growth rate *µ*_max_ was a good approximation of the intrinsic growth rate *µ*. We calculated the relative fitness of the mutant with respect to the wild-type as 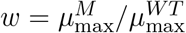 (or the selection coefficient as 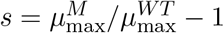).

Figure 4A shows that using *µ*_max_ to estimate the relative fitness generally infers incorrect values for the relative fitness (as well as the selection coefficient), unless both strains have exactly the same initial population fraction (*N*_0_*/K*). Even more concerning, this estimation sometimes categorizes the mutant as deleterious when it is beneficial, and vice versa. This is especially worrisome because we assumed noise-free data and an ideal case in which both bacterial strains follow the same population dynamics. In summary, equivocating *µ*_max_ with *µ* is likely to lead to wrong estimates of fitness and selection coefficients.

**Figure 4:**
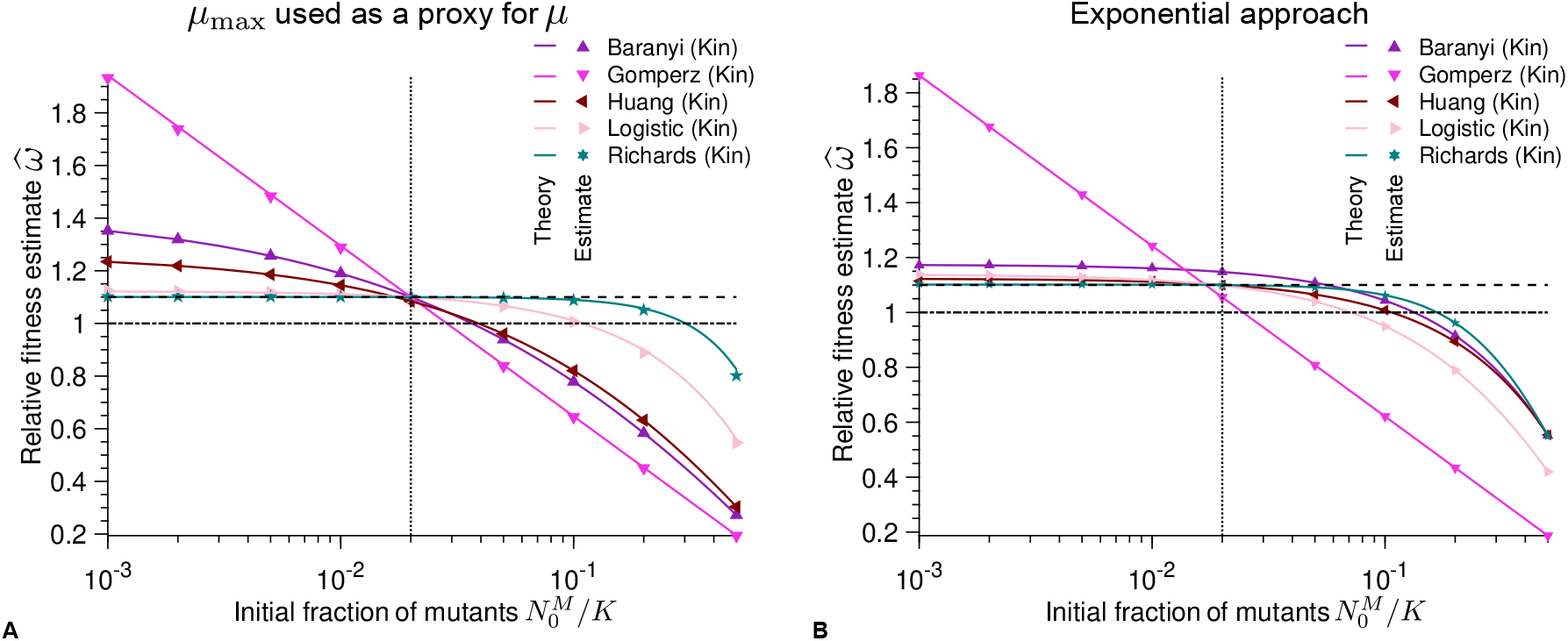
Different initial population fractions between wild-type and mutant batch cultures result in poor estimates of relative fitness: Relative fitness 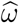 estimate versus initial fraction *N*_0,*M*_ */K* of mutants for different kinetic population growth models. As a reminder, *ω* = *µ*_*M*_ */µ*_*WT*_. **A) the maximum growth rate** *µ*_max_ **is used as a proxy for the intrinsic growth rate** *µ*. Each point represents estimated values by Spline (growthrates package) from simulated data averaged over 10^4^ stochastic realizations. The solid lines correspond to the analytical predictions of the relative fitness using estimates of the maximum growth rate (see Table 2). **B) the intrinsic growth rate is obtained applying the exponential approximation**. Solid lines correspond to the analytical predictions of the relative fitness using estimates of the maximum growth rate (see Table 3). In both panels the dashed line shows the real relative fitness value *w*. The dotted line represents the configuration in which the growth model parameters of the mutant are equal to the parameters of the wild-type (except their intrinsic growth rates). The dash-dotted line corresponds to the neutral case, i.e. when both the mutant and the wild-type have the same growth rate. Parameter values: *K*_*W T*_ = *K*_*M*_ = *K* = 10^5^, *µ*_*W T*_ = 1, *µ*_*M*_ = 1.1, *α*_*W T*_ = *α*_*M*_ = 2, *β*_*W T*_ = *β*_*M*_ = 2, *h*_0,*W T*_ = *h*_0,*M*_ = 2, *τ*_*W T*_ = *τ*_*M*_ = 2 and *t* = 1.

Mis-estimation of the selection coefficient also occurs when calculating relative growth rate values using the exponential approximation. Lenski et al. (1991) extended the exponential approximation to calculate the fitness of a mutant strain (*M*) relative to the fitness of a wild-type strain 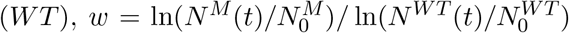 (or the selection coefficient 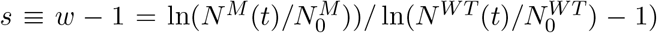. Note that under the assumption of exponential growth, both the relative fitness and the selection coefficient measured by the equation stated above are time-independent. Both Lenski et al. (1991) and Ram et al. (2019) empirically set the time interval of measurements *t* to 24 hours.

Similarly to using *µ*_max_ as a proxy for *µ*, the exponential approximation yields wrong estimates for the relative fitness (as well as the selection coefficient) in many cases (Figure 4B). For growth curves (except for the Gompertz model) in which the initial population fraction of the mutant is smaller than that of the wild-type, the exponential approximation is a more conservative estimator than *µ*_max_; it is less likely to overestimate the relative fitness. However, when the initial population fraction of the mutant is larger than that of the wild-type, the exponential approximation is likely to incorrectly infer that a beneficial mutant is deleterious (Figure 4B). Thus, the exponential approximation both misestimates and misclassifies throughout much of the experimentally reasonable parameter range.

##### Implications for the estimation of selection coefficients from growth rates

We showed that using *µ*_max_ as a proxy for *µ* in order to calculate the relative growth rates or relative fitness can lead to biased estimates (Figure 4). The amount of bias in the estimate becomes larger as the difference in the true growth parameters between the two studied strains becomes larger. Accordingly, we strongly recommend that experimenters use the intrinsic growth rate *µ* to estimate relative fitness. According to population biology, relative fitness is defined as the ratio of the intrinsic growth rate of the mutant strain over the wild-type strain, *µ*^*M*^ */µ*^*WT*^ (Lenski et al., 1991; Crow and Kimura, 2009; Chevin, 2011). From this point of view, it follows that relative fitness can only be estimated using the intrinsic growth rate *µ*. Nevertheless, fitness-related phenotypes, like 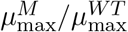, are sometimes used as a summary statistic of population growth that is contextual to the environmental, temporal, and population conditions (e.g., Adkar et al. 2017). Above we showed that a proxy of the intrinsic growth rate (like *µ*_max_ or the exponential approximation) can accurately estimate the relative fitness only if specific criteria are met. When these criteria are *not* met, *µ*_max_ becomes a composite parameter that depends on other experimental quantities that should be reported, like the initial fraction and lag time. Only if experiments are set up with small and equal initial fractions of the mutant and wild-type strains and if the strains do not exhibit a lag phase, using *µ*_max_ as a proxy for *µ* to estimate the relative fitness may be justified.

### 2.4 Application of theory: Re-analyzing 4 published data-sets

In order to clarify the theoretical considerations discussed above, we re-analyzed four published data-sets using the diversity of methods discussed above and then compared how the different methods performed on the same data. Among the 50 studies reviewed, we identified four that were appropriate for re-analysis since these papers provided their complete data-set and reported their estimated values (Adkar et al., 2017; Hammer et al., 2021; Ram et al., 2019; Todd and Selmecki, 2020). Each of these bacterial growth curve data-sets reports optical density versus time for different bacterial strains, with a total of 142 curves across all studies. We used three publicly available R packages, Growthcurver, grofit and growthrates, to estimate growth parameters (Sprouffske and Wagner, 2016; Kahm et al., 2010; Petzoldt, 2020). We tested two model-free methods (Spline and Easy Linear) and the model-based methods listed in Table 1. These models are based on different equations that are statistical (Zwietering et al., 1990) or kinetic (Tsoularis and Wallace, 2002; Baranyi and Roberts, 1994; Huang, 2011). We focused on inferring the maximum growth rate (*µ*_max_), both because it is of greatest relevance to the work discussed above and because it is the only quantity common to all methods we tested. For kinetic models (which are defined in terms of *µ*), we estimated *µ*_max_ by using derivatives to find the maximum slope of ln(*N/N*_0_) (as shown in Figure 1C; see Table 2).

The maximum growth rate (*µ*_max_) values estimated from the same data vary widely depending on the method used (Figure 5A, and Figures S1A, S2A and S3A). It is not possible to assess the accuracy of the estimates for the model-free methods. However, we used goodness-of-fit tests to assess the model-based methods by calculating the residual sum of squares (RSS). The RSS measures the discrepancy between the data and the fitted model. Thus, the smaller the RSS, the better the model. Since the models we tested have different numbers of parameters, we also calculated Akaike’s Information Criterion (AIC) for kinetic models using the method of López et al. (2004). The results of the AIC are consistent with those of the RSS (see Supplementary Material). Despite the discrepancy in the inferred maximum growth rate, many of the models fit the data well in most cases because the RSS values are low and similar (Figures 5B-C, and Figures S1B-C, S2B-C and S3B-C).

**Figure 5:**
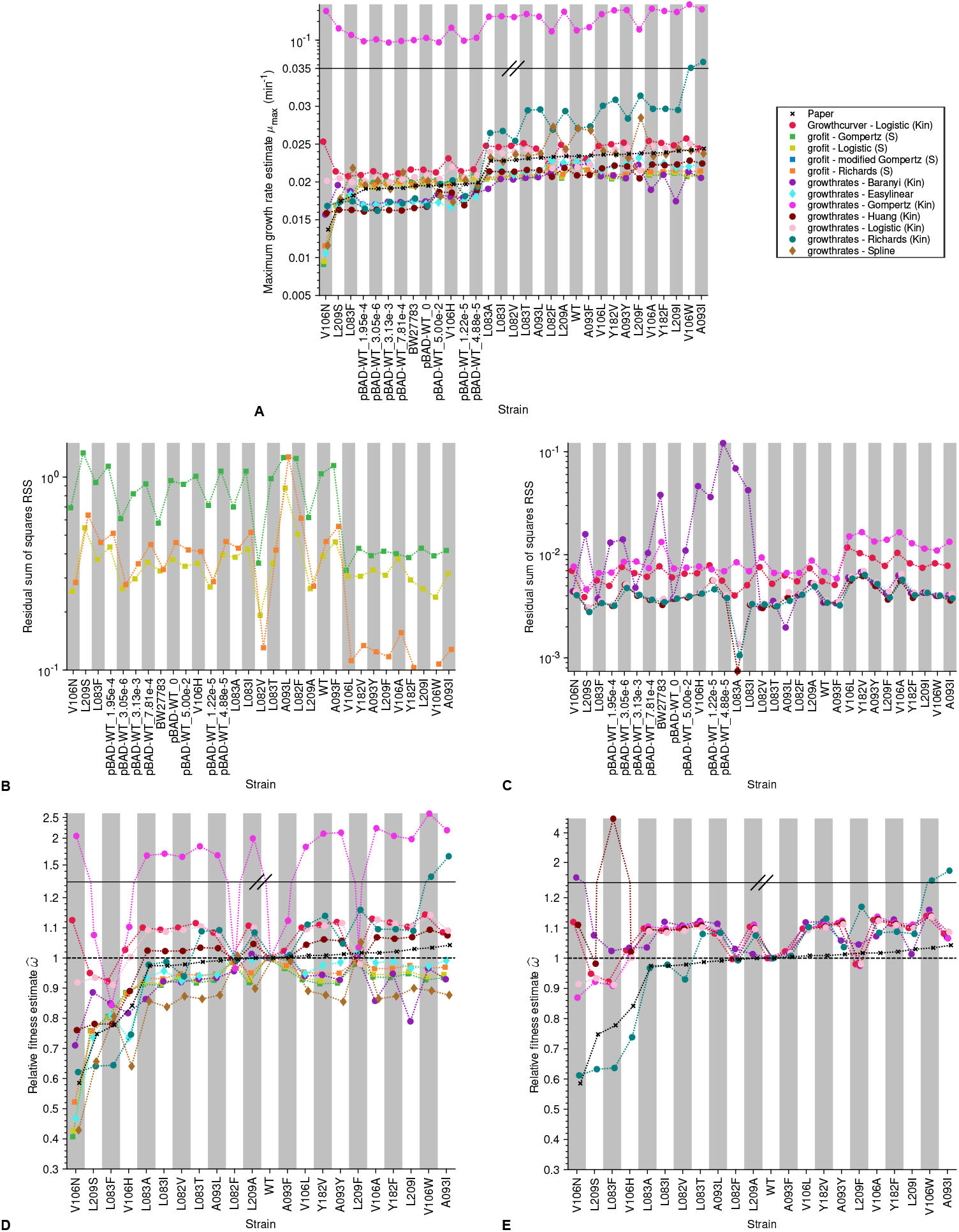
Analysis of published data-sets (Adkar et al., 2017) shows that estimates differ vastly depending on the method: A) Maximum growth rate 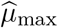 estimate for each strain. Each growth curve was analyzed using three different R packages including both model-free and model-based methods. The crosses show the values reported in the paper, the circles are obtained by methods based on kinetic models, the squares by methods based on statistical models and the diamonds by model-free methods. B) Goodness-of-fit measured as residual sum of squares (RSS) by strain for the statistical models. C) RSS by strain for the kinetic models. We include the RSS of statistical and kinetic models in different plots as the scale of the y-axes differ between these models: statistical models operate on a logarithmic scale since *y* = ln(*N/N*_0_) but kinetic models operate on a linear scale *N* (*t*). **Relative fitness:** and E) Relative fitness estimate 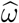 versus strain. In D) the relative fitness is computed using *µ*_max_ whereas in E) it is computed using *µ*.

A visual inspection of the fits corroborates our findings that all models fit the data generally well (visualizations of all fits available at available at https://github.com/LcMrc/GrowthRates). We emphasize that a visual inspection is important to ensure that the estimated values are appropriate. Indeed, in a few cases, the fits proved to be unsatisfactory although the summary statistics were good (see, e.g., Figure S6).

No model is consistently preferred for all samples of a data-set. This highlights the difficulty of choosing ‘the one’ right model, although this choice has a great impact on the growth parameter estimates. The Gompertz equation, whether statistical or kinetic, frequently yielded the worst statistics, although our literature review indicates the statistical Gompertz equation as the most frequently used model. Moreover, the models used in the original publications of the data were not always the models that we found to obtain the best statistics.

As previously stated, model-free methods may be preferred for some research questions, especially when neither the underlying mechanisms nor relative fitness estimates are of interest. Model-free methods obtain the maximum growth rate by determining the maximum of the function d ln(*N* (*t*)*/N*_0_)*/*d*t* (see Figure 1C). A quick comparison with the derivative of the experimental data ensures the validity of the estimate. We found that Easy Linear gave slightly different results from the Spline method because the former requires the user to specify how many data points to include for the analysis of the log-linear part of the growth curve. Note that model-free methods are likely to be more accurate than model-based methods for estimating *µ*_max_ because the latter involve more parameters and a data fit.

#### Relative growth rate estimates from empirical data

Adkar et al. (2017) used *µ*_max_ estimates from a statistical Gompertz model in order to estimate relative fitnesses. Therefore, this study allowed us to evaluate our concerns about using *µ*_max_ for relative fitness estimates. First, we used all approaches (model-free methods, statistical models, and kinetic models) to estimate *µ*_max_ for the wild-type and mutant strains from this data-set. Assuming (erroneously) that *µ*_max_ was a proxy for *µ*, we then calculated the relative fitness. Figure 5D shows that kinetic models (circles) estimate more beneficial fitness values than methods that estimate *µ*_max_ directly (diamonds for model-free methods and squares for statistical models). Especially for strains that were estimated by Adkar et al. 2017 (crosses) to have especially low fitness, we found a large variation in the fitness values estimated by different methods.

Next, we used kinetic models to estimate the intrinsic growth rate *µ* for the wild-type and mutant strains from this data-set and subsequently calculated the relative fitness. In Figure 5E we compared our estimated values with those published by Adkar et al. (2017). Again, many models estimate larger fitness values than those reported in the original study. This is especially pronounced for samples that were estimated by Adkar et al. 2017 (crosses) to have especially low fitness.

The overall conclusion of this section is that estimating relative fitness using inaccurate estimates of *µ* likely propagates to the level of relative fitnesses, and causes large discrepancies between relative fitness values estimated using different methods. Importantly, these discrepancies are most pronounced for samples that are of special interest in an experiment.

#### Implications of data re-analysis

Our analysis indicates that the choice of the best method to analyze growth curve data is very difficult. Especially, identifying the ‘right’ model that best fits all strains/treatments within a data-set seems daunting. This difficulty might explain why there exists such a diversity of methods for analyzing growth curve data, as we found in the literature review. Interestingly, we saw that most articles only report using one method for analyzing their data. We suspect that different labs and researchers have their own preferences and habits on how to obtain growth rate estimates. In the interest of time (and sanity), researchers may be using the model and method they know best and for which they have obtained reasonable-looking results in the past, rather than trying out a multitude of unfamiliar computational tools.

One clear finding from our re-analysis of published data is that the Gompertz family of models – both statistical and kinetic – are usually not the best choice. This point was previously made by López et al. (2004). We corroborate their empirical results with mathematical arguments. However, our literature review showed that the Gompertz models have remained popular long after López et al.’s study in 2004. We recommend that experimenters fit and compare more than one model when analyzing data. In the light of our results, it will be important to develop one easy-to-use framework that allows for model choice and comparison, which would easily single out inappropriate models.

Our results confirmed the finding of (Peleg and Corradini, 2011) that standard statistical techniques for model selection were often unhelpful (Figure 5B-C): when comparing the fit of different models to the same data, the goodness-of-fit statistics did not always select the model that looked best upon visual inspection and/or there was not much difference between models in terms of goodness-of-fit. This corroborates our personal communications with empiricists that they are reluctant to use models for fitting their growth curve data.

## 3 Recommendations & Conclusions

Despite the long-established study of batch culture growth curve data, estimating growth rates is still not straightforward. Using a literature review, math, simulations, and analysis of previously published data, our work highlights experimental and theoretical pitfalls encountered by many researchers who work with batch monoculture growth curves. To that end, we have summarized our recommendations for better growth rate estimates as a checklist in Figure 6.

**Figure 6:**
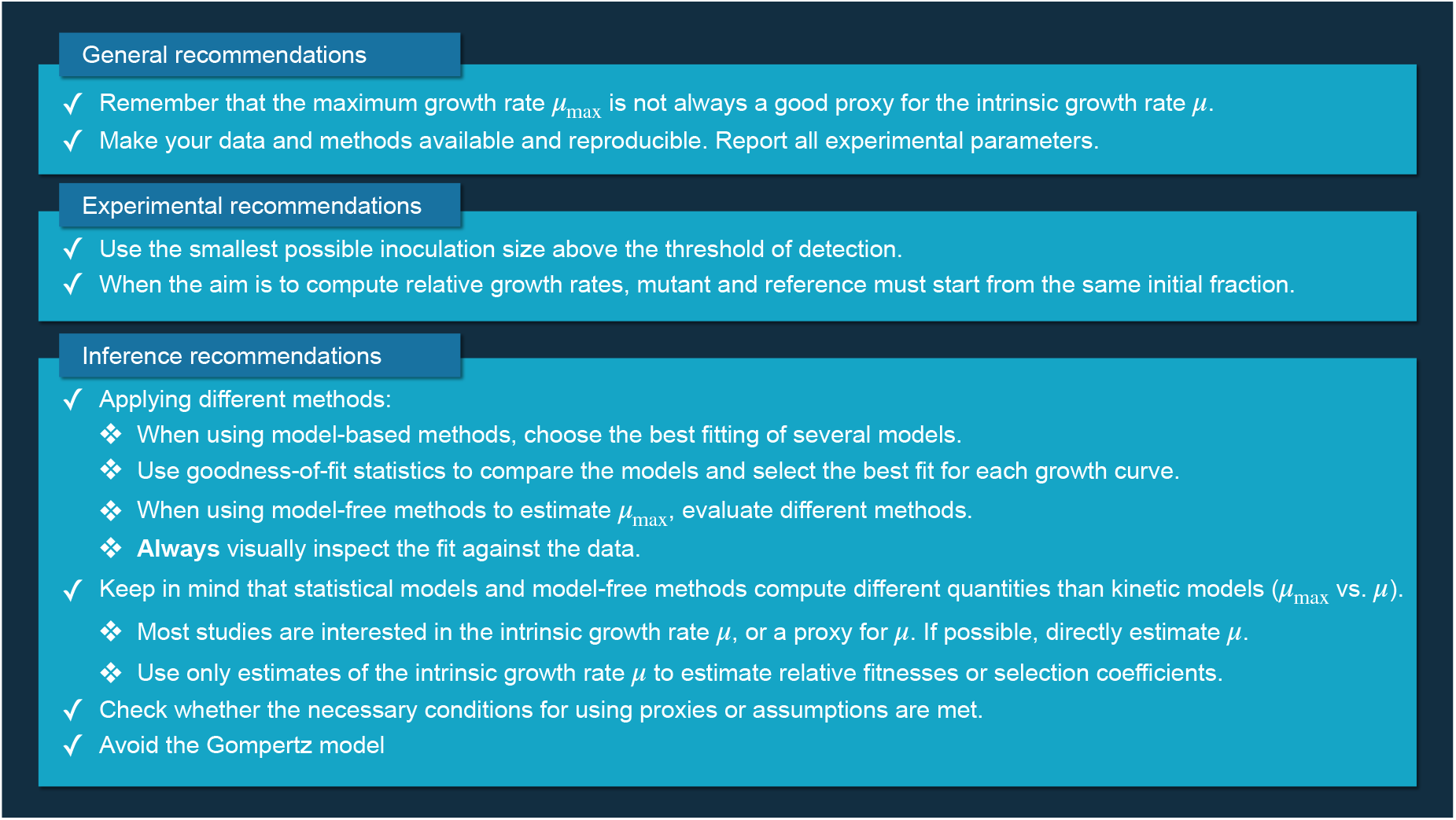
A list of recommendations for better growth rate inference from monoculture growth curves.

### General recommendations

We urge readers to remember that the intrinsic growth rate *µ*, a model parameter estimated from kinetic models, is not the same as the maximum growth rate *µ*_max_, a statistical quantity of data estimated using statistical models or by model-free methods. Although *µ*_max_ is often used as a proxy for *µ*, this assumption is not always justified.

We recommend that researchers make their raw data and methods available and reproducible. In particular, this involves reporting all experimental parameters like inoculum size, carrying capacity or initial fraction, and lag time. During our literature review, we were surprised by the lack of sufficient information on experimental methods, estimated values, and data availability.

### Experimental recommendations

Good data begin with good experimental methods. In the absence of a lag phase, the fastest growth rate (both for the maximum growth rate *µ*_max_ and for the intrinsic growth rate *µ*) is observed at the very start of the growth curve. As we show in Figures 3A and 4, the estimates of interest can be highly sensitive to the inoculum size. Also, when a lag phase is suspected, reliable data from the start of the growth curve are necessary to quantify the lag time and growth rate. Therefore, we recommend that experimenters use the smallest inoculation size possible that is still reliably above the threshold of detection (if using a microplate reader, follow the recommendations of Hall et al. 2014).

For accurate estimation of the relative growth rate, we stress that the mutant and reference/wildtype strains should start from the same initial fraction 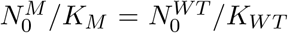; Figure 4), which is not necessarily the same absolute size. If the strains have the same carrying capacity (*K*_*M*_ = *K*_*WT*_) at stationary phase, then it is possible to either dilute the cultures used for inoculation by the same dilution factor or begin the growth curves at the same absolute inoculum size 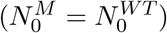. However, if the carrying capacities differ between strains, then the same absolute inoculum size cannot be used. We found in our literature review that about half of papers use a fixed absolute inoculum size to start their growth curve experiments whereas the other half use a fixed dilution factor, usually without justifying either choice. We recommend that experimenters interested in calculating relative growth rates use a fixed dilution factor. A pilot experiment is ideal to inform on the carrying capacities and corresponding optimal inoculum sizes.

### Inference recommendations

Regarding inference methods, our main recommendation is to try out several different methods on the same data. We recommend that a computational method is used to estimate the growth rate. (Manual) Fitting of an exponential model should be avoided (Table 3), regardless of whether the exponential model is fit explicitly or implicitly by using the equation *R* = (log_2_ *N*_2_ − log_2_ *N*_1_)*/*(*t*_2_ − *t*_1_).

Model-free computational methods tend to be technically easier to use than model-based methods, but they can only estimate *µ*_max_. If a model-free method is used, we recommend to try different programs, ideally based on different algorithms. Model-based methods often require more computational knowledge for fitting, but we still recommend that researchers try to fit more than one model. We recommend this because, by re-analyzing previously published data, we showed that there is not one model that fits best for all data-sets or even one model that best fits every curve within a specific data-set (Figures 5B-C, S1B-C, S2B-C and S3B-C). Therefore, it is important to select the best fitting model for each curve. For those familiar with the R statistical programming language, we recommend the growthrates package, because all of the kinetic models presented in Table 1, as well as model-free methods for estimating *µ*_max_, can easily be fitted with this package. We recommend that researchers use goodness-of-fit summary statistics (like the RSS, AIC, etc.) to compare models and select the best fit. However, additional visual inspection of the estimated value and the data is essential because goodness-of-fit statistics can be misleading (e.g., Figure S6).

Our work explains and demonstrates that and why the maximum growth rate *µ*_max_ is different from the intrinsic growth rate *µ*, which is a key point to take away and remember from this paper. We suggest that researchers attempt to estimate *µ* directly from their data using kinetic models. However, we have shown that it can be justified to use the statistical quantity *µ*_max_ as a proxy for the model parameter *µ* if certain conditions are met. We recommend that experimenters decide which one makes more sense to use for the experimental question at hand and, based on that decision, select the types of models or model-free approaches to use. If *µ*_max_ is to be inferred, model-free methods or fits of statistical models can be used. Model-based methods can either be used to estimate the maximum growth rate *µ*_max_ of statistical models, or the intrinsic growth rate *µ* of kinetic models. We urge experimenters not to compare estimates obtained from kinetic and statistical models because these different model types estimate different growth rate parameters. Researchers should be aware that confusion between the two quantities is common and different authors/fields use different naming conventions. We hope that this paper provides readers with the necessary conceptual understanding to critically navigate the literature. Finally, we note that only the intrinsic growth rate *µ* (and not *µ*_max_ or the exponential approximation) should be used for estimating the relative fitness (Figure 4) from monoculture growth curves.

It is important to verify that the necessary conditions are met for the inference method(s) to be used. For example, the exponential approximation should only be used when the following assumptions are met: the inoculum size is much smaller than the carrying capacity (*N*_0_ « *K*, e.g., by at least 2 orders of magnitude), only time points at which the population size remains much smaller than the carrying capacity are considered (*N* (*t*) « *K*), and the lag time is very short or absent (e.g., *h*_0_ « 1 for growth following a Baranyi model). Given these restrictive assumptions, rather than demonstrating that the conditions for the exponential approximation are met, it may be more feasible for researchers to directly fit one of the kinetic models listed in Table 1. Another approximation that requires justification is the use of *µ*_max_ as a proxy for *µ*. This approximation is only valid for small initial population fractions (Figure 3) and short (or absent) lag times. To demonstrate that this is the case, it is essential that experimenters report the initial population fraction(s) and the lag time(s) when using *µ*_max_ as a proxy for *µ*.

Finally, we recommend that researchers avoid fitting the Gompertz model for both absolute and relative estimates of *µ*. In our study, the kinetic Gompertz model consistently showed wrong estimates for simulated data (Figures 3-4) and unusually large estimates for empirical data (Figure 5).

## 4 Methods

### 4.1 Literature review

We quantified the most frequently used methods of analyzing growth curve data for extracting summary statistics. To this end, we used Web of Knowledge and Google Scholar to search for peer-reviewed papers in the evolution and ecology literature that gathered any type of growth curve data proportional to the number of individuals growing in a homogeneous, liquid culture and estimated growth parameters from those data. Papers that quantified binary presence/absence of growth (e.g. to assay lag time or spore viability) were excluded. Most papers were found because they cited one of the following growth curve methods papers, Zwietering et al. (1990); Hall et al. (2014); Sprouffske and Wagner (2016); or Delaney et al. (2013). From each paper, we extracted information about whether the method used to analyze growth curves is explicitly cited or described, what type(s) of growth curve summary statistic was used, whether a model-free or model-based approach was used, whether the growth rate is from a statistical or kinetic approach, which model(s) were fitted (if no equation is given, then the name of the model as reported by the author), whether the growth curves were inoculated from a fixed starting value or as a fraction of the carrying capacity, and whether the growth curve raw data are publicly available or, at least, plotted (summarized in table S1). For all papers in which growth curves were inoculated using a fraction from overnight cultures, the dilution factor was used as an estimator of the initial dilution fraction. If given, we extracted the initial dilution factor and any accompanying information about the inoculum (e.g., length of overnight culture) to indicate whether the dilution factor is a good proxy for the initial population fraction (*N*_0_*/K*). For papers in which growth curves were inoculated using a fixed absolute number of cells, we report the dilution fraction only if sufficient information about the inoculum size and carrying capacity was provided in the methods. Finally, we categorized different model-free growth rate estimation methods that were applied *ad hoc* as either “Easy Linear” if a consistent method was given for selecting which points to include in the regression (since this is the main feature of popular model-free methods like that of Hall et al. 2014), or as “exponential approximation” if there was no information about which points were included in the regression or as “spline” if pairs of successive measurements were used to estimate the local slope of the curve.

### 4.2 Analyzing 4 published data-sets

Data-sets appropriate for our analysis were found during our literature review and the data were accessed as indicated in each paper.

We used the following R (version 4.1.1) packages to re-analyze the data: Growthcurver (version 0.3.1), grofit (version 1.1.1-1) and growthrates (version 0.8.2). Each of them was downloaded from the CRAN repository except grofit that we obtained from Kahm et al. (2010). Indeed the latter was found to be removed from the CRAN repository. The package Growthcurver is based on the kinetic logistic model, whereas grofit includes four statistical models (Logistic, Gompertz, modified Gompertz and Richards). The package growthrates provide both model-free methods (Easy Linear and Spline) as well as methods based on kinetic models (Logistic, Gompertz, Richards, Baranyi and Huang).

We analyzed 143 population growth curves (31 from Adkar et al. 2017; 6 from Ram et al. 2019; 66 from Todd and Selmecki 2020; and 40 from Hammer et al. 2021) using all methods mentioned above. We focused on the maximum growth rate *µ*_max_, because it is the only quantity common to all models and methods. Since the kinetic models are defined based on the intrinsic growth rate *µ*, we used Table 1 to calculate the maximum growth rate *µ*_max_ from the respective model.

To test the accuracy of the fits obtained by the model-based methods, we calculated the residual sum of squares (RSS). We used the definition from López et al. (2004):

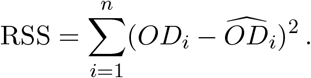

Here, *n* is the number of data points, *OD*_*i*_ is the *i*^th^ optical density value to be estimated and 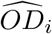 is the *i*^th^ estimated optical density value. Since the models have different numbers of parameters, we also calculated the Akaike’s Information Criterion (AIC) for the kinetic models as given in López et al. (2004) and explained in the supplementary methods.

### 4.3 Simulations

We generated data representing the dynamics of microbial populations using a Gillespie algorithm for the kinetic Gompertz, Richards and Logistic models (Gillespie, 1976, 1977). For the Baranyi and Huang models, a modified Next-Reaction algorithm was required since these models have time-dependent growth rates (Anderson, 2007). All simulation code was written in C and is available at https://github.com/LcMrc/GrowthRates. We detail below the algorithms used.

#### Gillespie algorithm

Let us denote by *N* the number of individuals. The only elementary event that can happen is division of a microbe, whose rate is denoted by *k*_*N*→*N*+1_. Let us note that *k*_*N*→*N*+1_ = *µ* log(*K/N*)*N, k*_*N*→*N*+1_ = *µ*(1 − *N/K*)*N* and *k*_*N*→*N*+1_ = *µ*(1 − (*N/K*)^*β*^)*N* for the kinetic Gompertz, Logistic and Richards models, respectively. Simulation steps are as follows:

1. Initialization: The population starts from *N*_0_ microorganisms at time *t* = 0.
2. Time update: The time increment Δ*t* is sampled randomly from an exponential distribution with mean 1*/k*_*N*→*N*+1_ and the time *t* is updated such that *t* ← *t* + Δ*t*.
3. Number of individuals update: a division occurs and the population size *N* increases by one such that *N* ← *N* + 1.
4. We go back to Step 2 and iterate until the desired time limit is reached.

#### Next-Reaction algorithm

Let us denote by *N* the number of individuals. The only elementary event that can happen is division of a microbe, whose time-dependent rate is denoted by *k*_*N*→*N*+1_(*t*). Let us note that 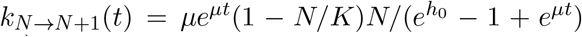 and *k*_*N*→*N*+1_(*t*) = *µ*(1 − *N/K*)*N/*(1 + *e*^*α*(*t*−*τ*)^) for the kinetic Baranyi and Huang models, respectively. In the following, we will denote by *P* the first firing time and *T* the internal time.

1. Initialization: The population starts from *N*_0_ microorganisms at time *t* = 0. The first firing time *P* is sampled from an exponential distribution of mean 1 and the internal time *T* is set to 0.
2. Time update: The time increment Δ*t* is computed solving 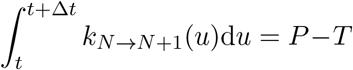 and the time *t* is updated such that *t* ← *t* + Δ*t*.
3. Number of individuals update: a division occurs and the population size *N* increases by one such that *N* ← *N* + 1.
4. Internal time update: The internal time *T* is updated such that *T* ← *T* + Δ*T*, where 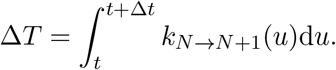.
5. First firing time update: The first firing time *P* is updated such that *P* ← *P* + Δ*P*, where Δ*P* is sampled from an exponential distribution of mean 1.
6. We go back to Step 2 and iterate until the desired time limit is reached.

### 4.4 Data availability

The authors state that all data necessary for confirming the conclusions presented in the article are represented fully within the article or Supplemental Material. Annotated C implementations of numerical simulations, annotated code to reproduce all computationally produced graphs, and additional figures and tables reporting the data re-analysis fits and estimates are available at https://github.com/LcMrc/GrowthRates.

## 5 Author Contributions

All authors designed the study; AHG performed the literature review; LM performed the numerical and analytical work; all authors analyzed and interpreted the data; all authors wrote and edited the manuscript.

## 6 Acknowledgements

The authors thank the Evolutionary Biology group at IGC for stimulating discussions and the THEE Group at UniBe for feedback on the manuscript. AHG thanks FCT PhD funding grant PD/BD/138215/2018. CB is grateful for funding from ERC Starting Grant 804569 (FIT2GO) and SNSF Project Grant “MiCo4Sys”.

## Supplement for

### 1 Supplementary methods

As explained in the main text, we re-analyzed the population growth curves from previously published data sets by fitting many different models and calculating the residual sum of squares (RSS). Since the models have different numbers of parameters, we also calculated the Akaike’s Information Criterion (AIC) for the kinetic models giving *N* (*t*), which reads

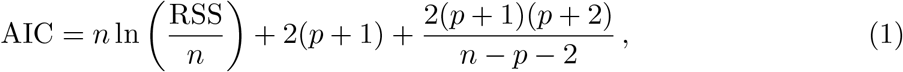

where *p* is the number of parameters (López et al. 2004).

However, this equation may not be valid for the statistical models because these models are on a logarithmic scale (*y* = ln(*N/N*_0_)) and therefore the errors around the data are probably not normally distributed as assumed by López et al. 2004.

### 2 Supplementary figures

**Figure S1:**
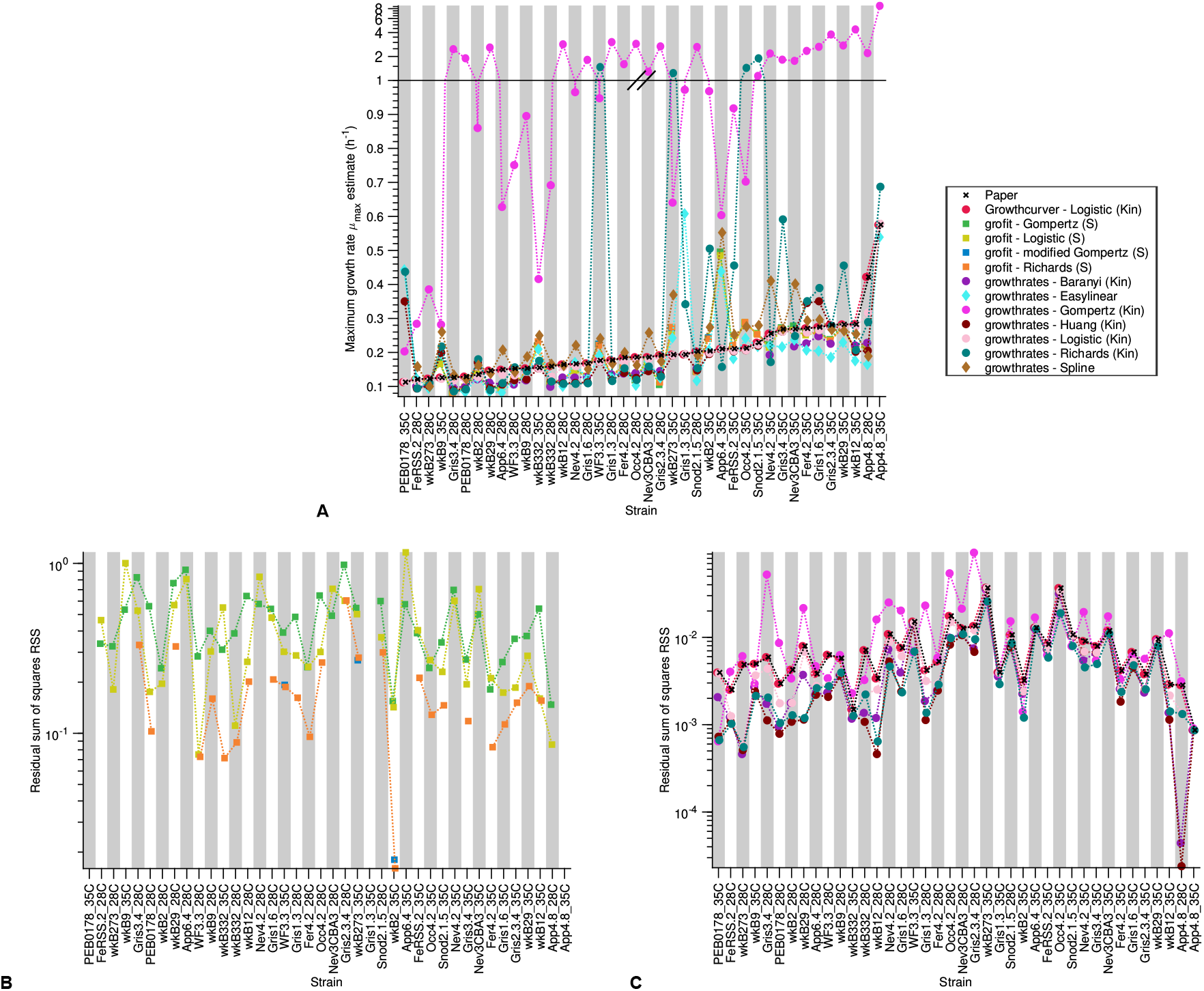
Analysis of published data sets - Hammer, Le, and Moran 2021: A) Maximum growth rate 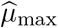 estimate versus strain. Each growth curve was analyzed using three different R packages including both model-free and model-based methods. The crosses show the values reported in the paper, the circles are obtained by methods based on kinetic models, the squares by methods based on statistical models and the diamonds by model-free methods. B) Residual sum of squares RSS versus strain for the statistical models. C) Residual sum of squares RSS versus strain for the kinetic models.

**Figure S2:**
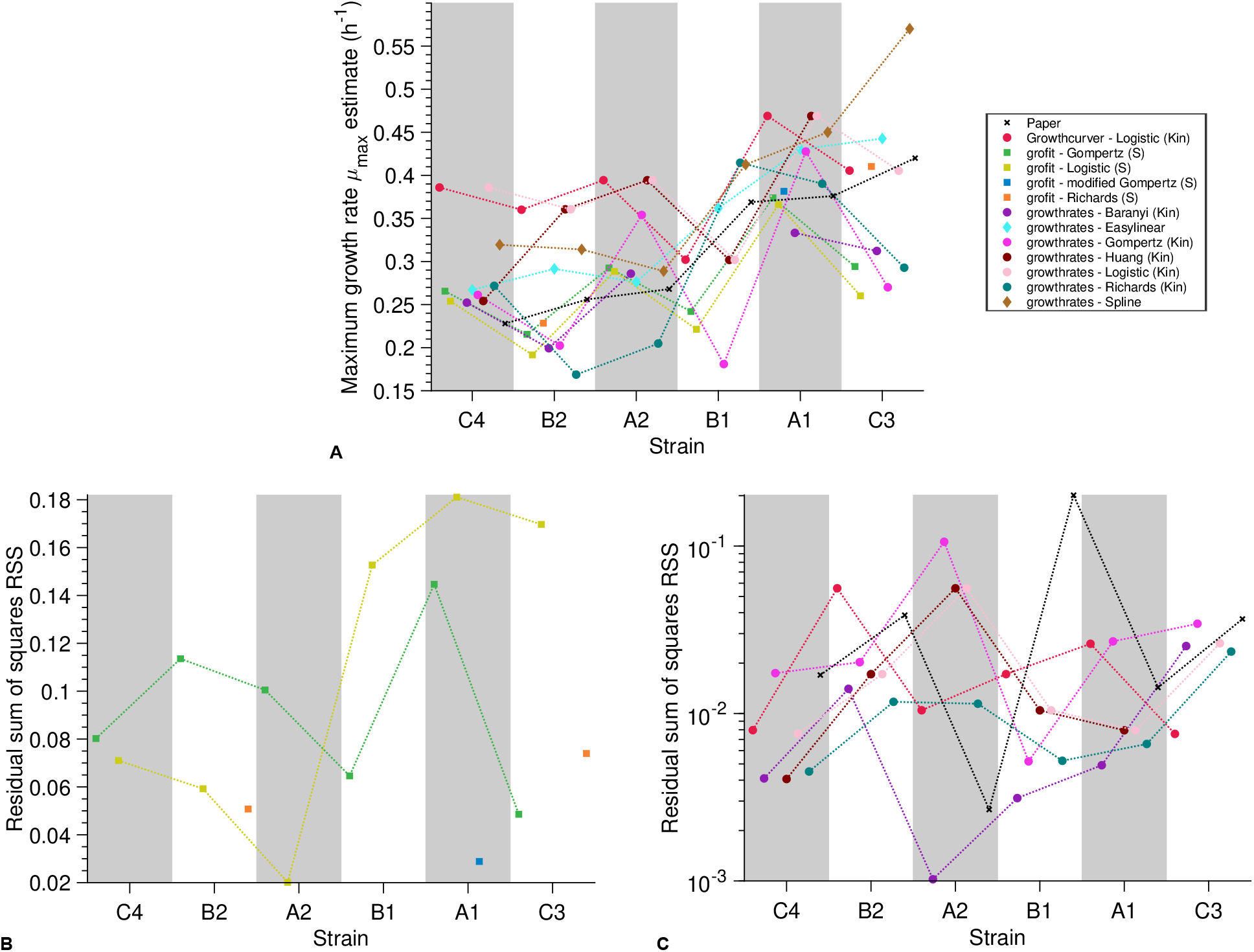
Analysis of published data sets - Ram2019: A) Maximum growth rate 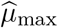 estimate versus strain. Each growth curve was analyzed using three different R packages including both model-free and model-based methods. The crosses show the values reported in the paper, the circles are obtained by methods based on kinetic models, the squares by methods based on statistical models and the diamonds by model-free methods. B) Residual sum of squares RSS versus strain for the statistical models. C) Residual sum of squares RSS versus strain for the kinetic models.

**Figure S3:**
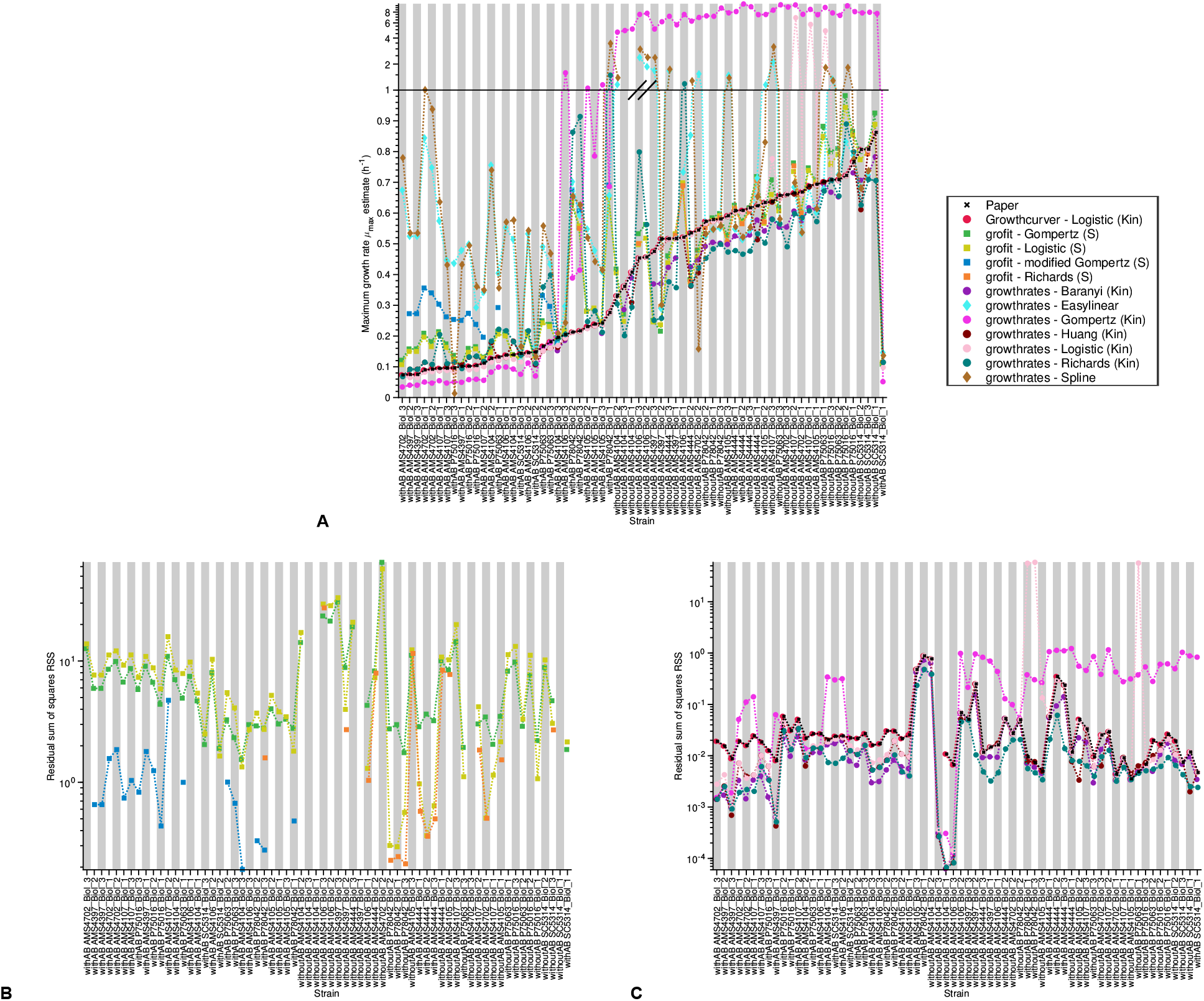
Analysis of published data sets - Todd and Selmecki 2020: A) Maximum growth rate 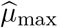 estimate versus strain. Each growth curve was analyzed using three different R packages including both model-free and model-based methods. The crosses show the values reported in the paper, the circles are obtained by methods based on kinetic models, the squares by methods based on statistical models and the diamonds by model-free methods. B) Residual sum of squares RSS versus strain for the statistical models. C) Residual sum of squares RSS versus strain for the kinetic models.

**Figure S4:**
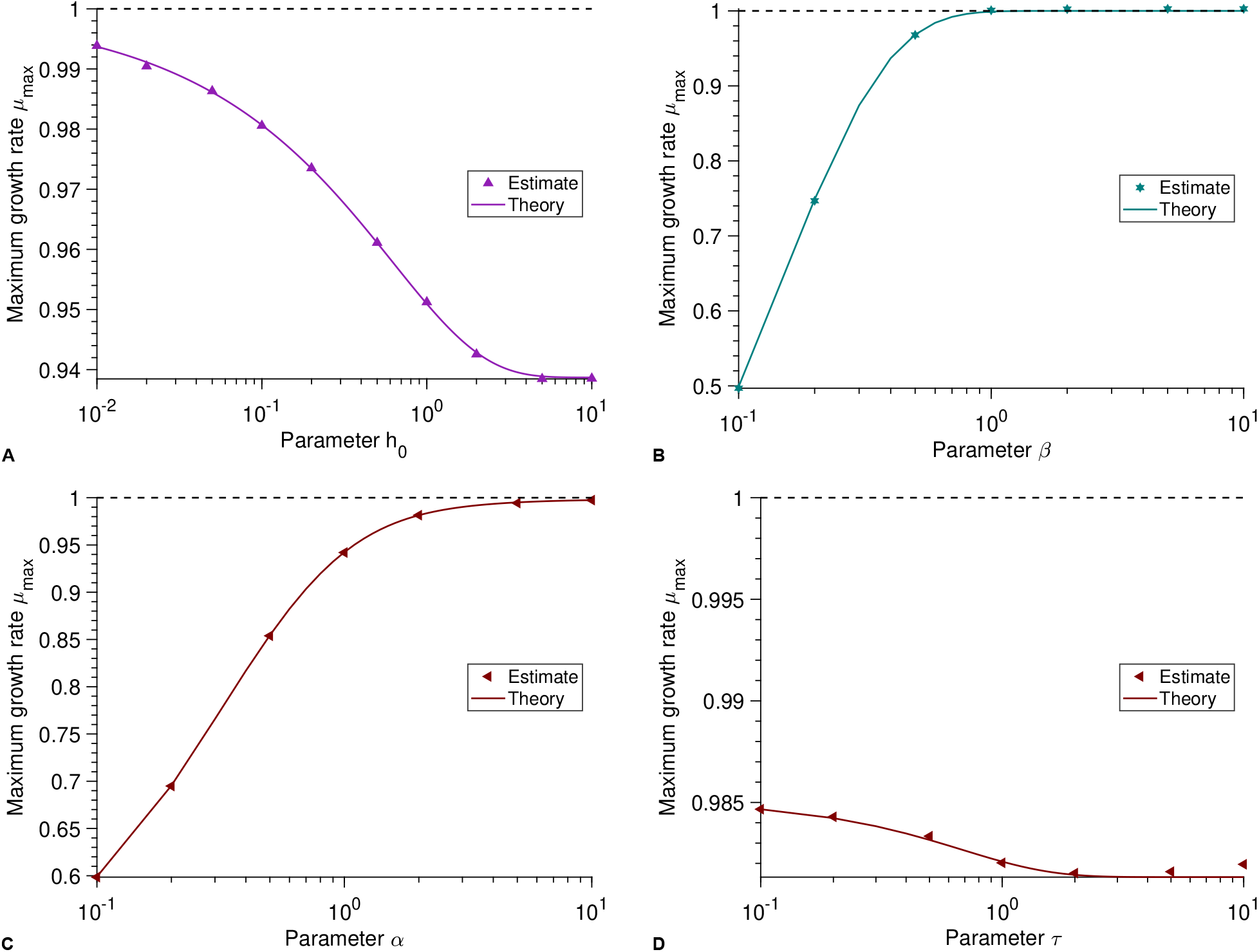
Maximum growth rate: A) Maximum growth rate *µ*_max_ versus parameter *h*_0_ for the Baranyi model. B) Maximum growth rate *µ*_max_ versus parameter *β* for the Richards model. C) Maximum growth rate *µ*_max_ versus parameter *α* for the Huang model. D) Maximum growth rate *µ*_max_ versus parameter *τ* for the Huang model. In every panel, each point represents estimated values from simulated data averaged over 10^4^ stochastic realizations. The solid lines correspond to the analytical predictions of the maximum growth rate (see Table 2 in the main text). The dashed line shows the intrinsic growth rate value *µ*. Parameter values: *N*_0_ = 10^2^, *K* = 10^5^, *µ* = 1, *α* = 2 for panel D and *τ* = 2 for panel C.

**Figure S5:**
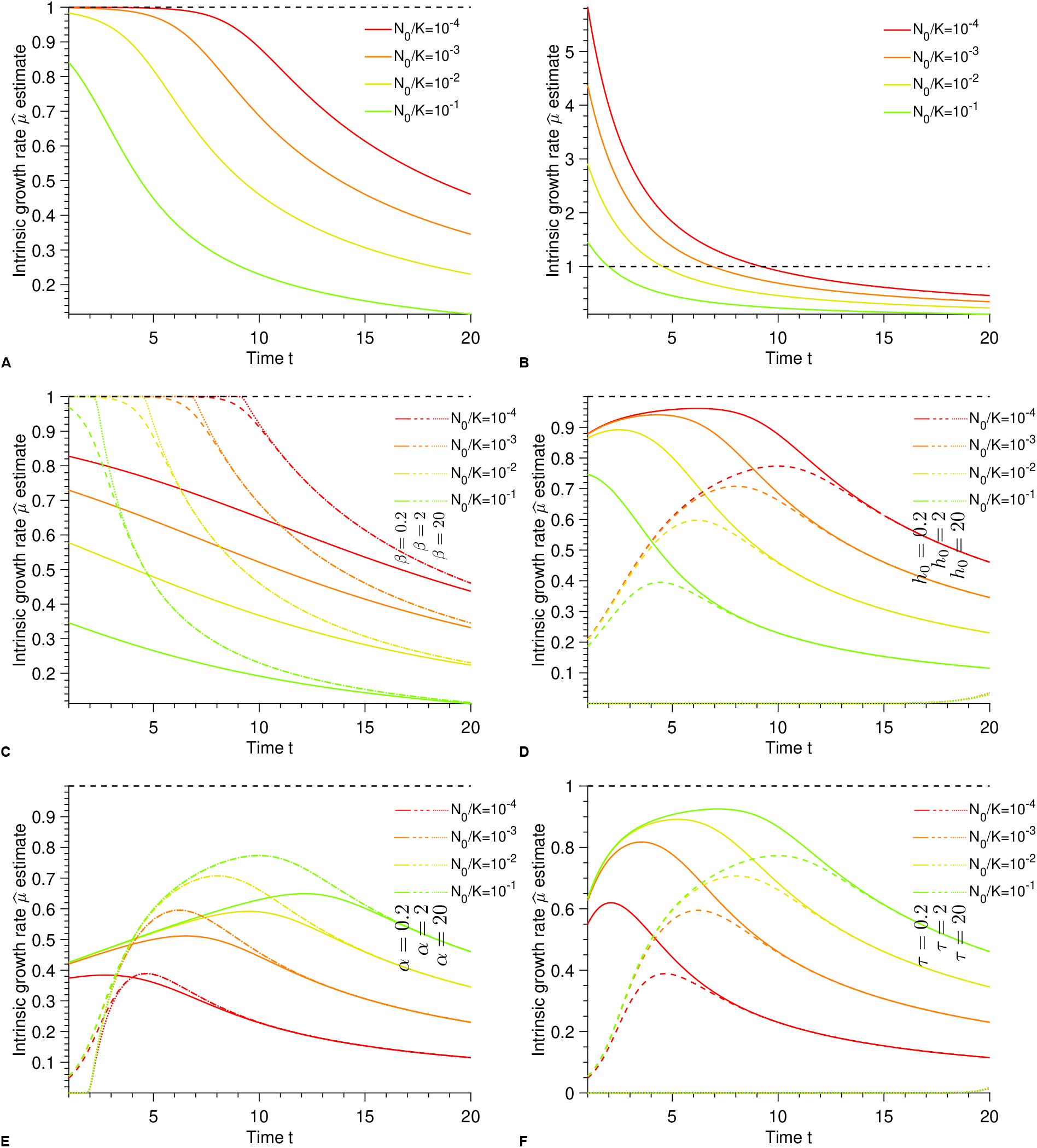
Intrinsic growth rate: From A) to F) Intrinsic growth rate 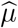 estimate versus parameter *t* for different initial population fractions *N*_0_*/K*, parameters and kinetic models (A) Logistic, B) Gompertz, C) Richards, D) Baranyi, E) and F) Huang). The intrinsic growth rate is analytically estimated using the exponential hypothesis *µ/* = ln(*N* (*t*)*/N*_0_)*/t*. The black dashed line shows the real value of *µ*. Parameter values: *K* = 10^5^, *µ, τ* = 2 for E) and *α* = 2 for F).

**Figure S6:**
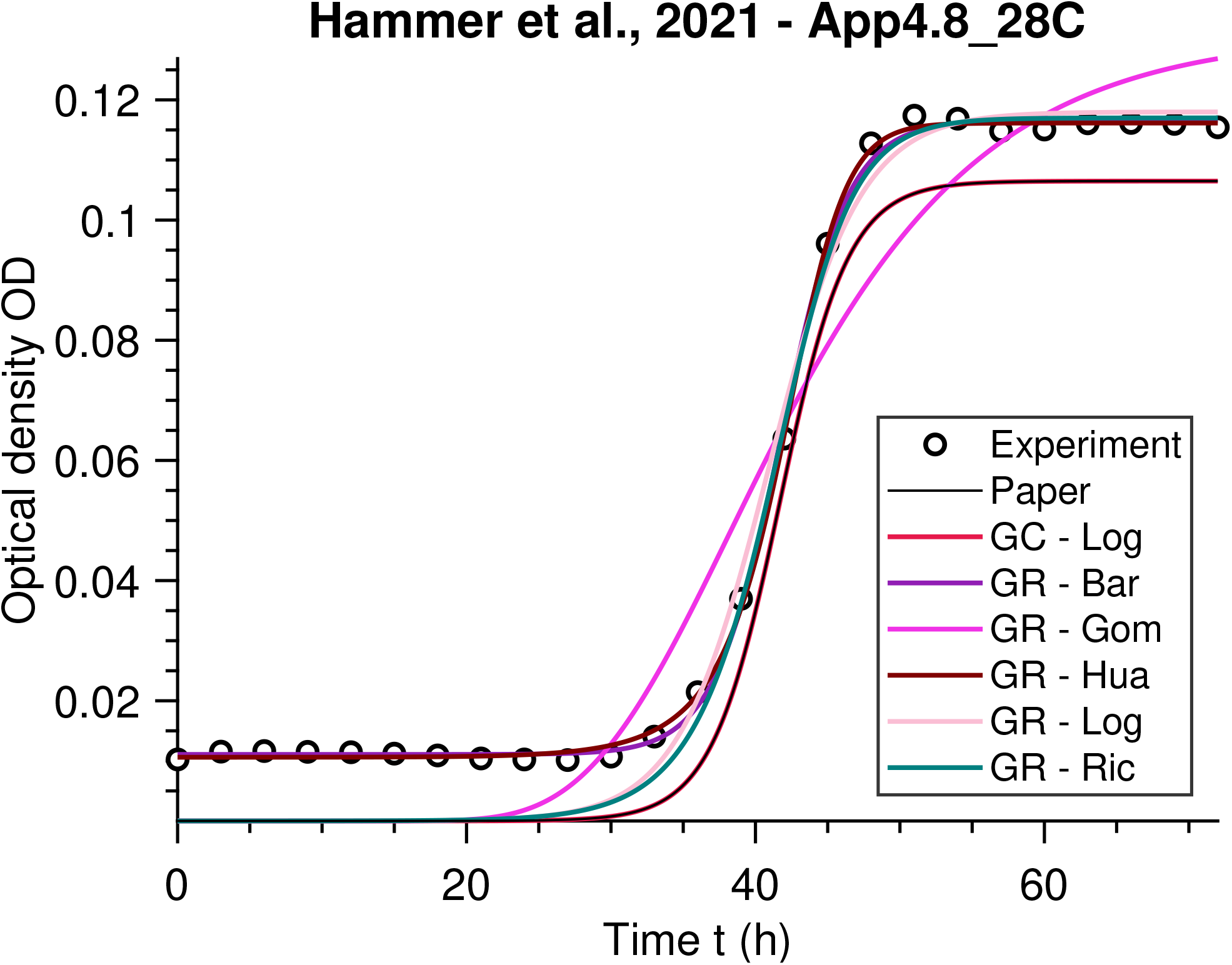
Analysis of published data: Optical density OD versus time *t*. The data points are fitted using different kinetic models. The black (Paper) and red (Growthcurver - Logistic (K)) lines are the same since it was the method applied in the paper.

**Figure S7:**
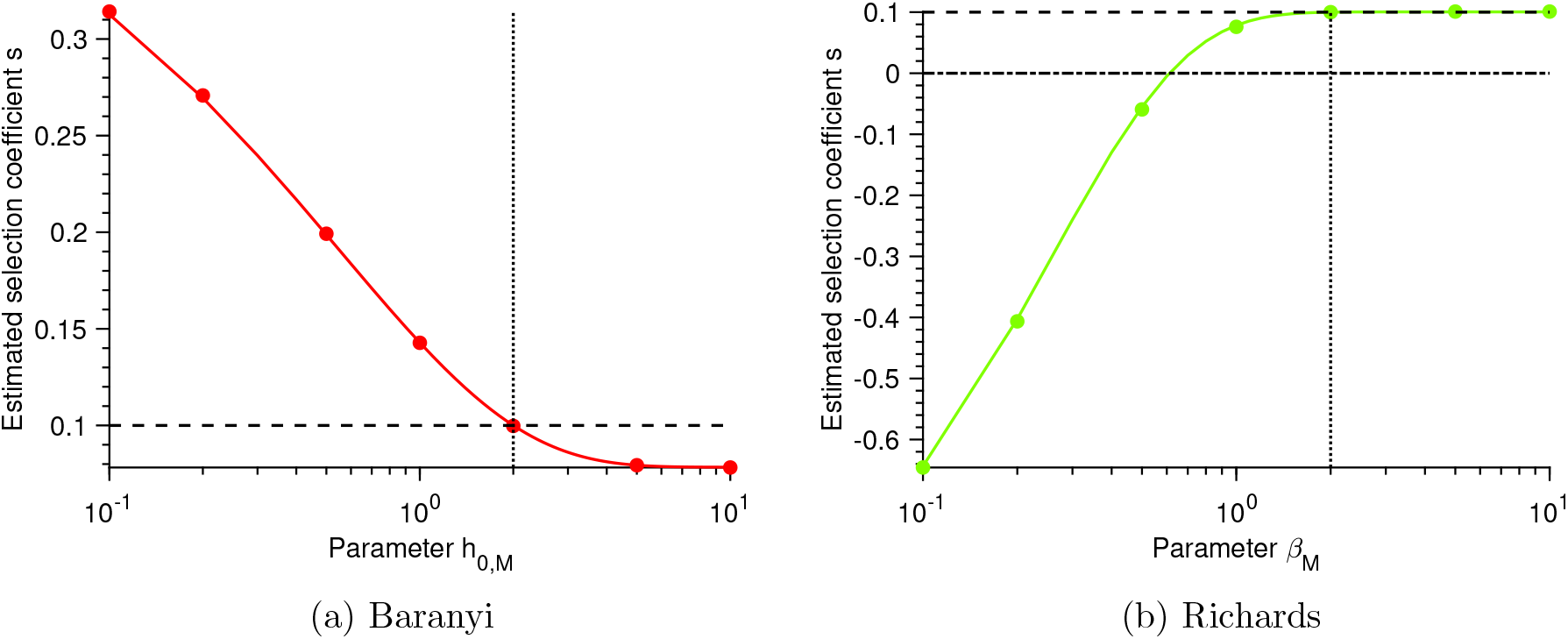
Selection coefficient: a) Estimated selection coefficient 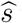 versus parameter *h*_0_ for the Baranyi model. b) Estimated selection coefficient 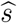 versus parameter *β* for the Richards model. In every panel, each point represents estimated values from simulated data averaged over 10^4^ stochastic realizations. The solid lines correspond to the analytical predictions of the selection coefficient using those of the maximum growth rate (see Table 2.2.1 in the main text). The dashed line shows the real selection coefficient value *s*. The dotted line represents the configuration where the parameters of the mutant are equal to those of the wild-type (except their growth rates). The dash-dot line corresponds to the neutral case, i.e. where both the mutant and the wild-type have the same growth rate. Parameter values: *K*_*W*_ = *K*_*M*_ = 10^5^, *µ*_*W*_ = 1, *µ*_*M*_ = 1.1, *s* = 0.1, *β*_*W*_ = 2 and *h*_0,*W*_ = 2. When not specified, *β*_*M*_ and *h*_0,*M*_ have the same values as the wild-type.

